# An essential role for miR-15/16 in Treg suppression and restriction of proliferation

**DOI:** 10.1101/2023.03.26.533356

**Authors:** Kristina Johansson, John D. Gagnon, Simon Zhou, Marlys S. Fassett, Andrew W. Schroeder, Robin Kageyama, Rodriel A. Bautista, Hewlett Pham, Prescott G. Woodruff, K. Mark Ansel

**Affiliations:** Department of Microbiology and Immunology, University of California, San Francisco, California, USA; Sandler Asthma Basic Research Center, University of California, San Francisco, California, USA; Department of Medicine, Division of Pulmonary and Critical Care Medicine, University of California, San Francisco, California, USA; Department of Medical Biochemistry and Cell Biology, University of Gothenburg, Gothenburg, Sweden; Department of Dermatology, University of California, San Francisco, California, USA; Department of Medicine, Genomics CoLab, University of California, San Francisco, California, USA

**Keywords:** T regulatory cell (Treg), miR-15, miR-16, immunosuppression, miRNA, effector Treg, TCF1

## Abstract

The miR-15/16 family is a highly expressed group of tumor suppressor miRNAs that target a large network of genes in T cells to restrict their cell cycle, memory formation and survival. Upon T cell activation, miR-15/16 are downregulated, allowing rapid expansion of differentiated effector T cells to mediate a sustained immune response. Here, using conditional deletion of miR-15/16 in immunosuppressive regulatory T cells (Tregs) that express FOXP3, we identify new functions of the miR-15/16 family in T cell immunity. miR-15/16 are indispensable to maintain peripheral tolerance by securing efficient suppression by a limited number of Tregs. miR-15/16-deficiency alters Treg expression of critical functional proteins including FOXP3, IL2Rα/CD25, CTLA4, PD-1 and IL7Rα/CD127, and results in accumulation of functionally impaired FOXP3^lo^CD25^lo^CD127^hi^ Tregs. Excessive proliferation in the absence of miR-15/16 inhibition of cell cycle programs shifts Treg diversity and produces an effector Treg phenotype characterized by low expression of TCF1, CD25 and CD62L, and high expression of CD44. These Tregs fail to control immune activation of CD4^+^ effector T cells, leading to spontaneous multi-organ inflammation and increased allergic airway inflammation in a mouse model of asthma. Together, our results demonstrate that miR-15/16 expression in Tregs is essential to maintain immune tolerance.

**Highlights:**

- Treg-specific miR-15/16 expression is essential to prevent systemic tissue inflammation
- miR-15/16 restrict Treg proliferation and regulate expression of the key functional Treg molecules FOXP3, IL2Rα, CTLA4, PD-1 and IL7Rα
- miR-15/16 limit formation of effector Tregs and is necessary for high suppressive capacity

## Introduction

Regulatory T cells (Treg) are essential mediators of peripheral immune tolerance. Severe functional Treg defects or complete lack of Tregs results in systemic immune activation and lethal autoimmunity (1–3). Treg function relies on IL-2 signaling via STAT5 that sustains high expression of FOXP3 and the high affinity IL-2 receptor α chain, CD25 (1,4–7). This positive feedback loop reinforces Treg transcriptional identity and promotes maturation of CD4^+^FOXP3^lo^ cells into CD4^+^FOXP3^+^CD25^hi^ Tregs during thymic development (8–10). More recent research has described functionally diverse Treg subgroups that depend on expression of specific transcription factors (11,12). For instance, gradient expression of TCF1 and LEF1 separates Tregs into subpopulations of resting and activated effector phenotypes with distinct transcriptional profiles (11). Functional specialization of Tregs play a critical role in preventing loss of self-tolerance (13), but the mechanisms that govern specification of Treg subgroups are incompletely understood.

MicroRNAs (miRNAs) are short non-coding RNAs that regulate gene expression by binding untranslated regions of target mRNAs to recruit Argonaute (Ago) proteins and promote mRNA degradation and translational repression (14). Coordinated regulation of gene networks gives miRNAs a profound biologic impact and it is well-known that miRNAs play crucial roles in differentiation and function of T cells (15–19).

The miR-15/16 miRNA family comprises four abundant miRNAs that are encoded in two separate clusters (miR-15a/miR-16-1 and miR-15b/miR-16-2). miR-15/16 have been studied extensively in lymphocyte malignancies, where they function as tumor suppressors (20,21). Deletion of miR-15a/miR-16-1 in the mouse *Dleu2/Mirc30* locus leads to clonal B cell lymphoproliferation that recapitulates many features of chronic lymphocytic leukemia (20). In more recent years, a role for miR-15/16 regulation of T effector responses has emerged (22–26), and one study reported that miR-15b/16-2 overexpression promoted FOXP3 expression and Treg differentiation (27). Yet our understanding of how miR-15/16 regulate Tregs remains limited.

Here, we generated mice with conditional inactivation of both miR-15/16 clusters in Tregs to determine their impact on Treg development and function. miR-15/16 restricted proliferation of thymus-derived Tregs and regulated the expression of proteins that are fundamental to Treg function, including securing high CD25 expression. Deletion of miR-15/16 in Tregs resulted in lymphoproliferative disease with systemic tissue inflammation and accumulation of Tregs with an activated effector phenotype.

## Results

### Treg-specific miR-15/16 expression is essential to prevent autoimmune inflammation

miR-15/16 are among the most abundant miRNAs occupying Ago2 complexes in resting T cells, but the miRNAs in this family are dynamically expressed. They are downregulated by T cell receptor (TCR) stimulation in vitro and remain low during memory T cell formation following lymphocytic choriomeningitis virus infection in mice (22). miR-15/16 expression in non-effector T cell subsets such as Tregs is less studied. We analyzed published miRNA expression data from sorted human peripheral blood T cells to assess miR-15/16 miRNAs in distinct effector and non-effector T cell subsets (28). Tregs exhibited the highest expression of miR-15a, −15b and −16 compared to CD4^+^ naïve, Th1, Th2 and Th17 cells (Figure S1).

To investigate the functional importance of miR-15/16 in Tregs specifically, we generated mice with conditional deficiency of miR-15 and miR16 using Cre recombinase driven by the endogenous *Foxp3* locus (*Foxp3*^Cre^). *LoxP*-flanked miR-15a/16-1 (*Mirc30*) and miR-15b/16-2 (*Mirc10*) alleles (hereafter referred to as miR-15/16^fl/fl^) were deleted in Tregs after Foxp3 was expressed. By the age of 20 weeks, miR-15/16^fl/fl^*Foxp3*^Cre^ mice developed spontaneous systemic inflammation and lymphoproliferation (Figure 1A-D). Specifically, we observed a significant infiltration of inflammatory cells in lung tissue, clusters of inflammatory cells in liver and thickening of the skin, but no apparent inflammatory features in the pancreas (Figure 1A). Furthermore, spleen and inguinal lymph nodes were visibly larger compared to those of miR-15/16^wt/wt^*Foxp3*^Cre^ and miR-15/16^fl/fl^*Foxp3*^Wt^ control mice (hereafter referred to as ‘WT control’) (Figure 1B), and spleens obtained from miR-15/16^fl/fl^*Foxp3*^Cre^ mice had significantly greater mass (Figure 1C). The total number of cells was increased in spleen and lymph nodes of miR-15/16^fl/fl^*Foxp3*^Cre^ mice compared to WT control mice (Figure 1D).

**Figure 1.**
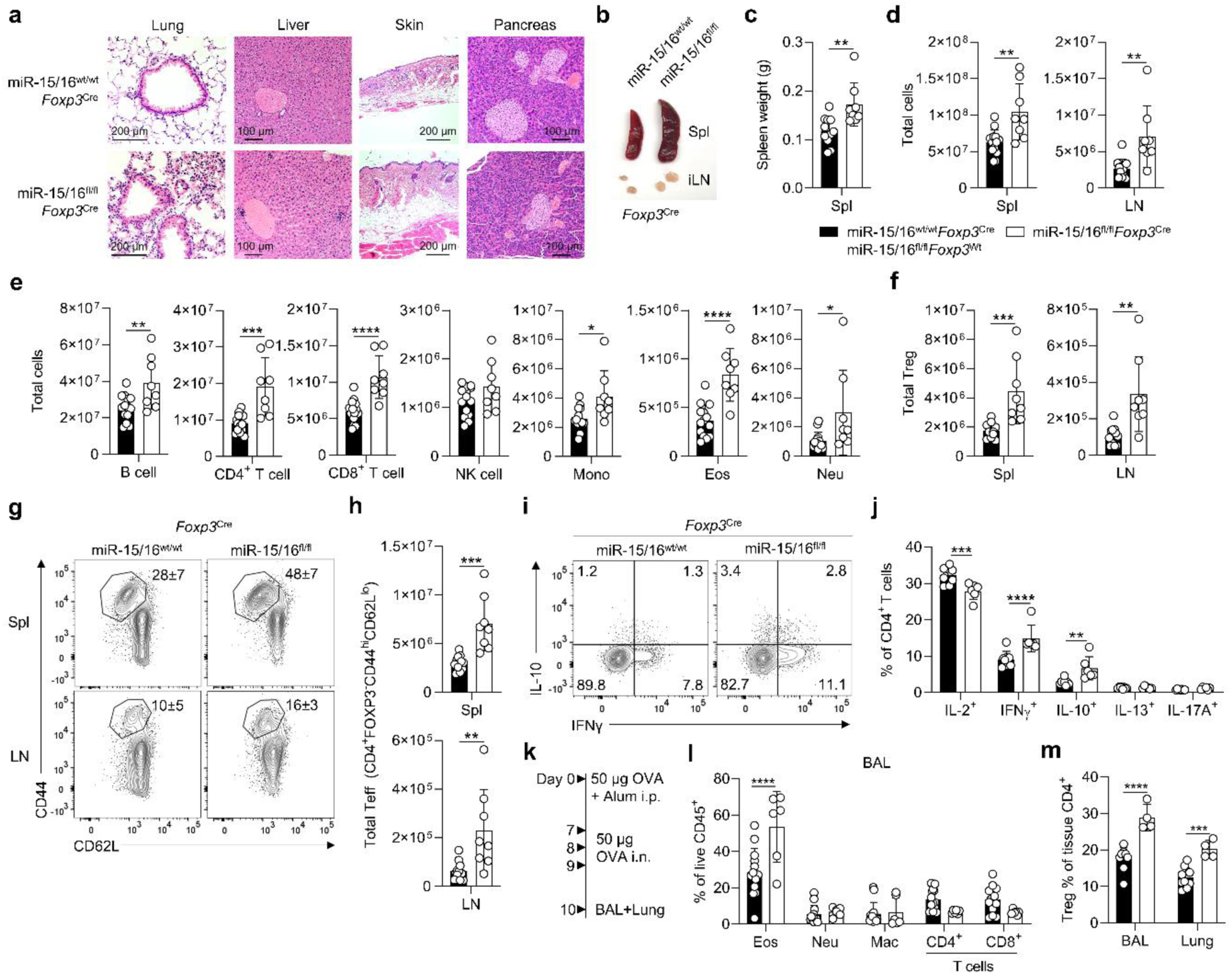
Treg-specific miR-15/16 expression is essential to prevent spontaneous and induced tissue inflammation. **A:** H&E-stained sections of lung, liver, skin and pancreas; control mice with miR-15/16 sufficient Tregs (miR-15/16^wt/wt^*Foxp3*^Cre^ or miR-15/16^fl/fl^*Foxp3*^Wt^; ‘WT’) on top row and mice with miR-15/16 deficient Tregs (miR-15/16^fl/fl^*Foxp3*^Cre^, ‘KO’) on bottom row. Representative image of spleen (Spl) and inguinal lymph nodes (LN) **(B)**, spleen weight quantified **(C)**, and total cells quantified in spleen and lymph nodes **(D)** from the same mice. Total spleen B cells (CD45^+^CD4^-^CD8^-^NK1.1^-^CD19^+^), CD4^+^ T cells (CD45^+^CD11b^-^CD11c^-^CD8^-^CD4^+^), CD8^+^ T cells (CD45^+^CD11b^-^CD11c^-^CD4^-^ CD8^+^), NK cells (CD45^+^CD4-CD8^-^CD19^-^NK1.1^+^), monocytes (Mono, CD45^+^CD11b^int-hi^CD11c^int-lo^NK1.1^-^Ly6G^-^), eosinophils (Eos, CD45^+^CD11b^+^Siglec-F^+^) and neutrophils (Neu, CD45^+^CD11b^+^Ly6G^+^) in spleen **(E)**, and total Tregs in spleen and lymph nodes **(F)** of WT and KO mice. Representative contour plots with frequencies **(G)** and quantification of total **(H)** CD44^hi^CD62L^lo^ T effector cells in spleen and lymph nodes of WT and KO mice. Intracellular cytokine staining of CD4^+^ T cells from WT and KO mice restimulated ex vivo; contour plots show IFN-γ and IL-10 expression **(I)** and quantification of IL-2^+^, IFN-γ^+^, IL-10^+^, IL-13^+^ and IL-17A^+^ cells **(J)**. **K:** Airway inflammation model induced by intraperitoneal (i.p.) OVA sensitization and intranasal (i.n.) OVA challenge. Bronchoalveolar lavage (BAL) and lung tissue were collected 24h after the last OVA challenge. Frequency of eosinophils (CD11b^+^Siglec-F^+^), neutrophils (CD11b^+^Ly6G^+^), alveolar macrophage (Mac, CD11c^+^CD11b^lo^), CD4^+^ T cells (CD4, CD11b^−^CD11c^−^CD4^+^) and CD8^+^ T cells (CD8, CD11b^−^CD11c^−^CD8^+^) among live hematopoietic cells in BAL **(L)**, and frequency of tissue-resident Tregs in BAL and lung tissue of OVA challenged WT and KO mice. 20-week-old unchallenged mice in A-J. Data from 7 independent experiments. N=6-14 mice/group. In C-F and H: Unpaired t-test 2-tailed and 2-way ANOVA with Bonferroni’s multiple comparison test in J and L-M. Bar graphs are shown with error bars demonstrating standard deviation.

Splenocytes analyzed by flow cytometry exhibited increased abundance of a variety of inflammatory cells including B cells, CD4^+^ and CD8^+^ T cells, monocytes, and eosinophil and neutrophil granulocytes (Figure 1E). Focusing on CD4^+^ T cells, miR-15/16 deficiency in the Treg compartment led to higher numbers of both Tregs (Figure 1F) and T effector (Teff) cells (Figure 1G-H), indicating that the absence of miR-15/16 in Tregs has secondary effects on the effector response. T cell cytokine production was analyzed by intracellular flow cytometry following PMA/Ionomycin stimulation ex vivo, revealing a higher frequency of IFN-γ- and IL-10-producing cells, but lower frequency of IL-2-producing cells, in miR-15/16^fl/fl^*Foxp3*^Cre^ mice compared to WT controls (Figure 1I-J). There were no differences in the proportion of IL-13- or IL-17-producing cells (Figure 1I-J). Taken together, our findings of systemic inflammation in multiple tissues and activation of effector cells suggest that miR-15/16 are essential for normal Treg functions that maintain immune homeostasis and prevent spontaneous inflammation.

### miR-15/16-expressing Tregs better suppress induced inflammation

To investigate the impact of miR-15/16 in Tregs under inflammatory conditions following an immunological trigger, we sensitized mice with a single intraperitoneal injection of ovalbumin (OVA) in alum (day 0) and challenged the airways with OVA on three consecutive days (day 7, 8, 9) one week later (Figure 1K). Before sacrificing the mice, a fluorophore-conjugated antibody to the hematopoietic marker CD45 was injected intravenously allowing separation of airway-resident cells from blood cells by flow cytometry. Bronchoalveolar lavage (BAL) collected 24h after the last OVA challenge showed significantly higher frequency of airway-infiltrating eosinophils in miR-15/16^fl/fl^*Foxp3*^Cre^ mice compared to WT controls (Figure 1L), despite similar numbers of neutrophils, macrophages, and CD4^+^ and CD8^+^ T cells (Figure 1L). The airway eosinophilia in miR-15/16^fl/fl^*Foxp3*^Cre^ mice was accompanied by an accumulation of airway-resident Tregs in the lung tissue and BAL (Figure 1M), indicating that miR-15/16 deficient Tregs are unable to suppress OVA-induced inflammation as effectively as WT Tregs.

### miR-15/16 selectively restrict Treg expansion

Our previous studies in *Cd4*-Cre mice, where miR-15/16 deficiency affects all T cells, demonstrate that miR-15/16 act through direct binding and posttranscriptional regulation of cell cycle-associated genes to limit the expansion of CD4^+^ and CD8^+^ T cells during antiviral responses (22), but possible variations within the CD4^+^ T cell compartment were not investigated. Like the miR-15/16^fl/fl^*Foxp3*^Cre^ mice, miR-15/16^fl/fl^ *Cd4*-Cre mice exhibited an overgrowth of Tregs (Figure 2A-B). In fact, the increase in Tregs was selective as there was no difference in the number of FOXP3^-^ conventional T cells (Tcon) in miR-15/16^fl/fl^ *Cd4*-Cre mice compared to miR-15/16^fl/fl^ controls (Figure 2B). To examine whether the accumulation of Tregs was associated with heightened proliferation, we injected intravenous 5-ethynyl-2’- deoxyuridine (EdU) into miR-15/16^fl/fl^ *Cd4*-Cre and control mice carrying a FOXP3-GFP reporter (Figure 2C-D). Among CD4^+^ T cells of the thymus, spleen and lymph nodes, we detected increased EdU-incorporation in GFP^+^ Tregs of miR-15/16^fl/fl^ *Cd4*-Cre mice, but not in GFP^−^ Tcons (Figure 2E). We distinguished distinct Treg subsets of likely thymic (tTreg) or peripheral (pTreg) origin by their expression of Helios or NRP-1, respectively (Figure 2F) (29). The increased Treg pool in miR-15/16^fl/fl^ *Cd4*-Cre mice consisted of tTregs, as there was no significant difference in the number of pTregs (Figure 2G). In summary, these findings suggest that miR-15/16 specifically restrict the accumulation of thymically-derived Tregs, but not at the expense of conventional T cells or peripherally induced Tregs.

**Figure 2.**
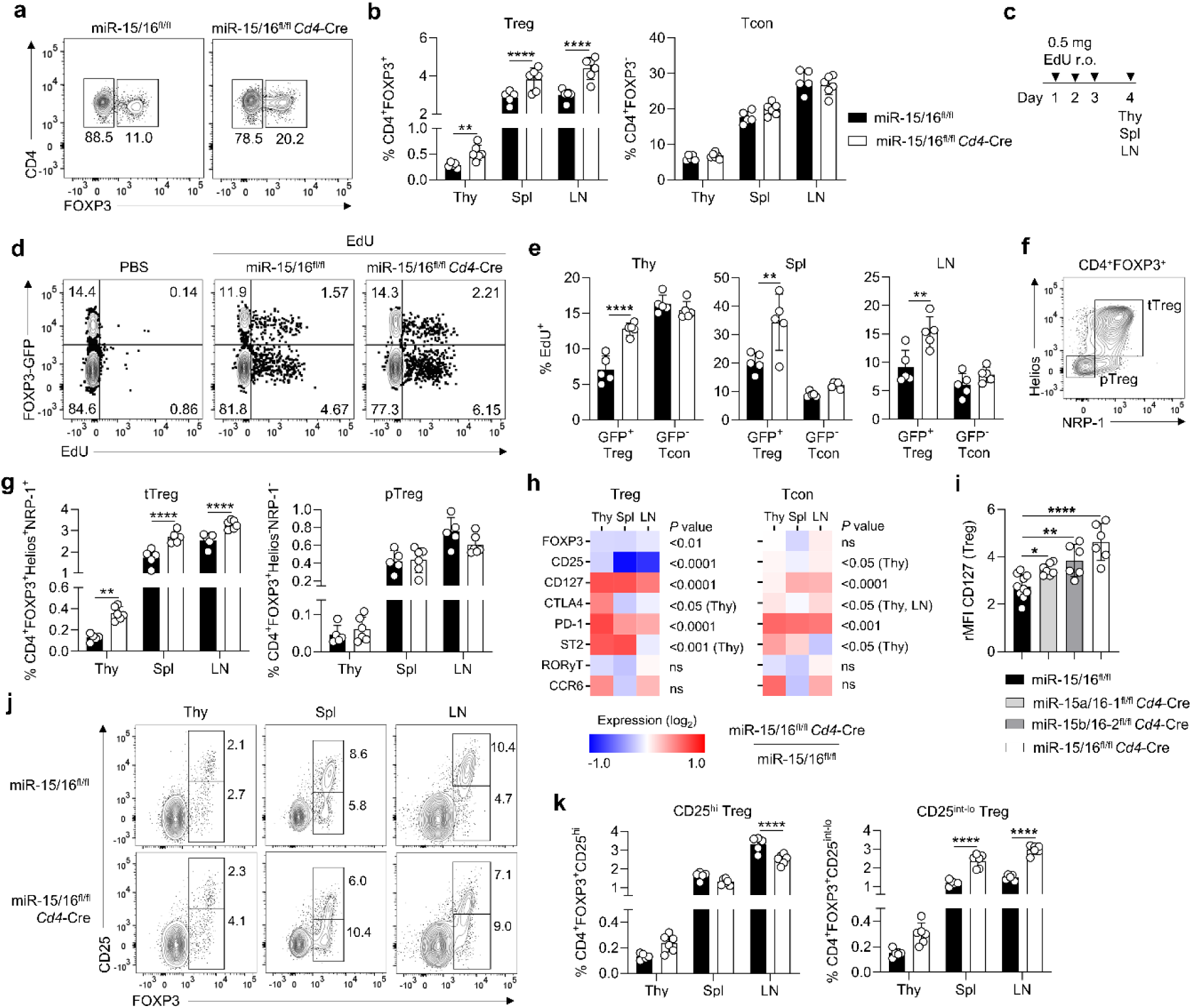
miR-15/16 specifically restrict the expansion of Tregs and regulate expression of key Treg proteins. Representative contour plots **(A)** and quantification of frequencies **(B)** of CD4^+^FOXP3^+^ Tregs and CD4^+^FOXP3^-^ conventional T cells (Tcon) among live hematopoietic cells in thymus (Thy), spleen (Spl) and lymph nodes (LN) of miR-15/16^fl/fl^ WT control and miR-15/16^fl/fl^ *Cd4*-Cre mice. **C:** Model to track in vivo cell proliferation accomplished by retroorbital (r.o.) injection of 5-ethynyl-2’-deoxyuridine (EdU). Tissues were collected 24h after the last injection. **D:** Representative contour plots of EdU-injected miR-15/16^fl/fl^ WT control, miR-15/16^fl/fl^ *Cd4*-Cre mice and PBS control mice all carrying a FOXP3-GFP reporter. **E:** Quantification of frequencies of CD4^+^GFP^+^ Tregs and CD4^+^GFP^-^ Tcons of all CD4^+^ cells in indicated tissues of the same mice. **F:** Contour plot demonstrating gating strategy of thymically-derived tTregs (Helios^+^NRP-1^+^) and peripherally induced pTregs (Helios^-^NRP-1^-^). **G:** Quantification of frequencies of tTregs and pTregs in indicated tissues among live hematopoietic cells of miR-15/16^fl/fl^ WT control and miR-15/16^fl/fl^ *Cd4*-Cre mice. **H:** Fold change of median fluorescent intensity (MFI) by flow cytometry of indicated proteins in Tregs and Tcons from three tissues (change in miR-15/16^fl/fl^ *Cd4*-Cre from miR-15/16^fl/fl^ WT control). **I:** Relative MFI (rMFI) of CD127 expression in Tregs from miR-15/16^fl/fl^ WT control, single cluster deficient miR-15a/16-1^fl/fl^ *Cd4*-Cre mice and miR-15b/16-2^fl/fl^ *Cd4*-Cre mice and double cluster deficient miR-15/16^fl/fl^ *Cd4*-Cre mice. Representative contour plots **(J)** and quantification of frequencies **(K)** of CD25^hi^ Tregs and CD25^lo^ Tregs among live hematopoietic cells in indicated tissues of miR-15/16^fl/fl^ WT control and miR-15/16^fl/fl^ *Cd4*-Cre mice. Data from a minimum of 2 independent experiments. N=5-10 mice/group. In all bar graphs except I, 2-way ANOVA with Sidiak’s multiple comparison test. In I ordinary ANOVA with Dunnett’s multiple comparison test. Bar graphs are shown with error bars demonstrating standard deviation. 2-way ANOVA with Bonferroni’s multiple comparison test in heatmaps in H.

### miR-15/16 regulate the expression of key Treg proteins

As evident in the contour plots (Figure 2A) demonstrating Treg populations in miR-15/16^fl/fl^ *Cd4*-Cre and miR-15/16^fl/fl^ control mice, Tregs that lack miR-15/16 display abnormally low FOXP3 expression. We used median fluorescent intensity (MFI) in our flow cytometric analysis to quantify the amount of FOXP3 and other proteins that are critical to Treg functions including CD25, CD127, CTLA4 and PD-1. Tregs and Tcons obtained from thymus, spleen and lymph nodes were analyzed (Figure 2H). FOXP3 and CD25 expression were substantially reduced in miR-15/16^fl/fl^ *Cd4*-Cre Tregs compared to miR-15/16^fl/fl^ control Tregs (Figure 2H). In contrast, expression of the inhibitory receptors CTLA4 and PD-1 were enhanced in miR-15/16^fl/fl^ *Cd4*-Cre Tregs (Figure 2H). Slight but significant effects were also seen in Tcons, where PD-1 expression was most clearly affected (Figure 2H). However, expression of proteins not directly implicated in bona fide Treg functions (ST2, RORγT and CCR6) exhibited no or less consistent changes (Figure 2H). Not surprisingly, the direct miR-15/16 target CD127 was derepressed in absence of miR-15/16 in both Tregs and Tcons (Figure 2H), and by using mice lacking one of the two miR-15/16 family clusters, we identified a dose-response of CD127 derepression (Figure 2I). Indeed, in a stepwise fashion related to the abundance of the miRNA family, Tregs from mice with deficiency of both miR-15/16 clusters exhibited significantly higher CD127 expression compared to miR-15/16 WT and single cluster deficient mice (i.e. miR-15a/16-1^fl/fl^ *Cd4*-Cre and miR-15b/16-2^fl/fl^ *Cd4*-Cre) (Figure 2I). Further analysis of the impact of each miR-15/16 cluster on Treg cell number (in vitro and in vivo) and Treg phenotype is provided in supplementary Figure S2. These results show that miR-15/16 deficient Tregs have significant alterations in the expression level of central functional proteins. FOXP3^lo^CD25^lo^ Tregs accumulated in all tissues that were investigated in miR-15/16^fl/fl^ *Cd4*-Cre mice (Figure 2J-K). Together, the data suggest that miR-15/16 play an important role, not only in controlling the size of the Treg pool, but also in generating and maintaining a competitive Treg phenotype under homeostatic conditions.

### CD25^lo^ Tregs protect mice from intestinal inflammation

To test the functional implications of miR-15/16 deficiency in Tregs, we gave T cell deficient recipient mice 400,000 naïve WT T cells together with 150,000 Tregs (CD4^+^CD45.1^-^FOXP3-GFP^+^) from either miR-15/16 sufficient mice (‘miR-15/16^fl/fl^’) or miR-15/16 deficient mice (‘miR-15/16^fl/fl^ *Cd4*-Cre’) (Figure 3A). A third group received 400,000 naïve WT T cells alone (CD4^+^CD45.1^+^CD25^-^CD44^lo^CD62L^hi^; ‘Naïve only’) (Figure 3A). Recipient mice were monitored for wasting disease by body mass and condition over 8 weeks. Mice that received naïve T cells without Treg co-transfer developed colitis with visible inflammation of the colon (Figure 3B) and decreased body weight (Figure 3C). However, mice that received Tregs, from either miR-15/16^fl/fl^ or miR-15/16^fl/fl^ *Cd4*-Cre mice, had no visible inflammation of the colon (Figure 3B) or lower body weight by the end of the model (Figure 3C). The protective effect by Tregs was reflected by significantly lower frequencies of Teff cells (CD4^+^CD44^hi^CD62L^lo^) in the colon, mesenteric lymph nodes and spleen (Figure 3D). T cells expressing T-bet (Figure 3E) and IFN-γ (Figure 3F) were similarly reduced. The spontaneous accumulation of CD25^lo^ Tregs, previously found in mice lacking miR-15/16 expression in all T cells (miR-15/16^fl/fl^ *Cd4*-Cre) or in Tregs only (miR-15/16^fl/fl^ *Foxp3*^Cre^), was replicated in recipient mice that obtained miR-15/16 deficient Tregs (Figure 3G-H). This result suggest that miR-15/16-dependent Treg expansion is not controlled by external factors, a question we address with greater detail in the next section. Frequencies of CD25^hi^ Tregs were found at the same level in miR-15/16^fl/fl^ and miR-15/16^fl/fl^ *Cd4*-Cre Treg-recipients (Figure 3G-H). Significantly reduced expression levels of FOXP3 and CD25, and enhanced expression of CD127 and PD-1 were characteristic of miR-15/16 deficient Treg (Figure 3I), as reported in previous experiments. Accumulation of these abnormal Tregs resulted in higher Treg-to-Teff ratio in recipient mice (Figure 3J), possibly suggesting that Tregs lacking miR-15/16 expression compensate for impaired suppressive capacity and provide protection from intestinal inflammation by increasing in number.

**Figure 3.**
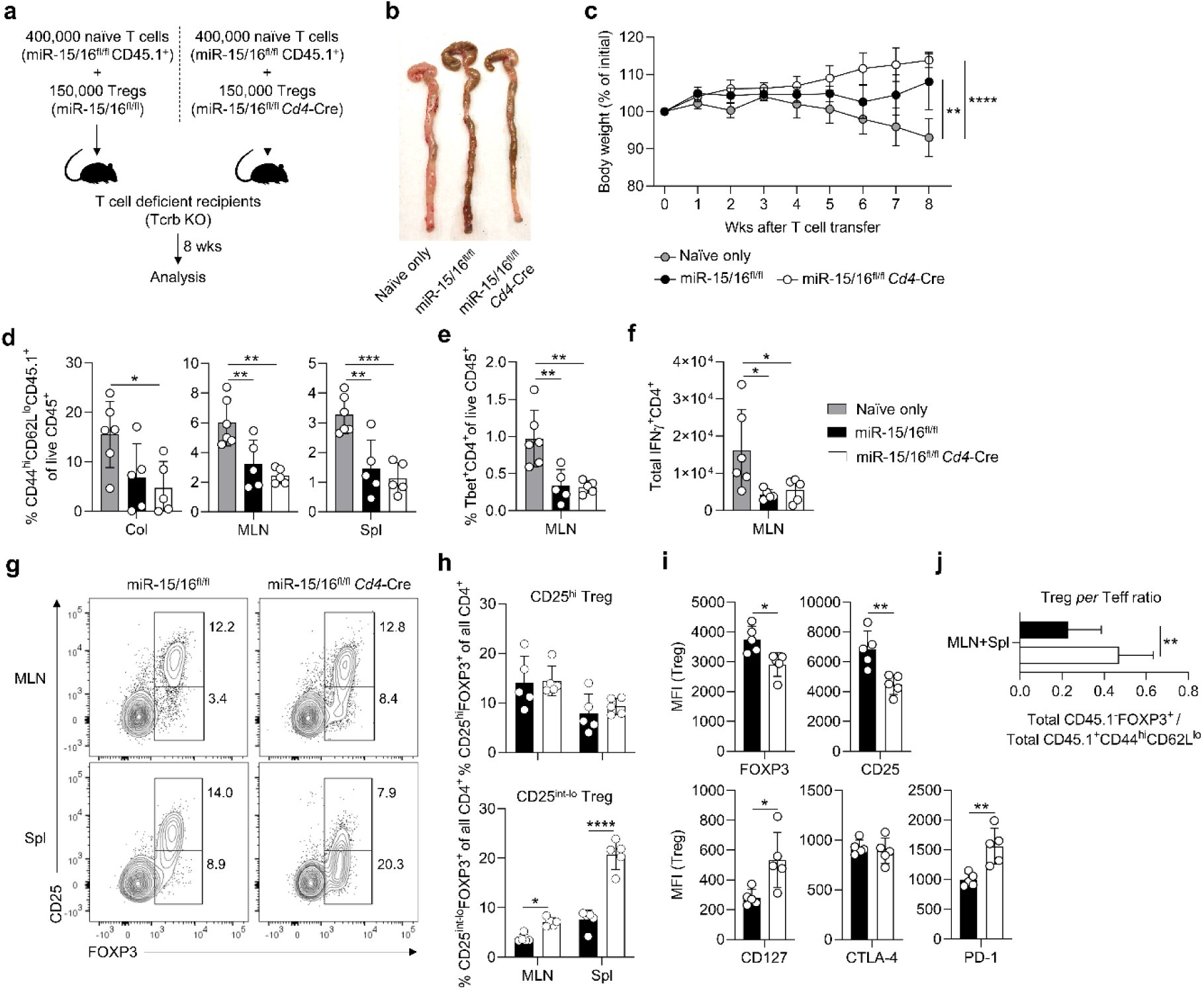
Accumulation of CD25^lo^ Tregs provides sufficient protection from intestinal inflammation. **A:** Colitis model by T cell transfer accomplished by intravenous injection of FACS-sorted naïve T cells and Tregs. Recipient mice were monitored for 8 weeks before analysis. **B:** Representative image of colons collected from mice the received naïve T cells (‘Naïve only’), naïve T cell together with miR-15/16 sufficient Tregs (‘miR-15/16^fl/fl^’), and naïve T cell together with miR-15/16 deficient Tregs (‘miR-15/16^fl/fl^ *Cd4*-Cre’). **C:** Body weight monitored weekly of the same mice. **D:** Quantification of frequencies of CD44^hi^CD62L^lo^ T effector cells among live hematopoietic cells in colon (Col), mesenteric lymph nodes (MLN) and spleen (Spl) of Naïve only, miR-15/16^fl/fl^ and miR-15/16^fl/fl^ *Cd4*-Cre recipient mice. Quantification of frequencies of Tbet^+^CD4^+^ T cells among live hematopoietic cells in MLN **(E),** and total IFN-γ^+^CD4^+^ T cells **(F)** in mesenteric lymph nodes of Naïve only, miR-15/16^fl/fl^ and miR-15/16^fl/fl^ *Cd4*-Cre recipient mice. Representative contour plots **(G)** and quantification of frequencies **(H)** of CD25^hi^ and CD25^lo^ Tregs among all CD4^+^ T cells in mesenteric lymph nodes and spleen of miR-15/16^fl/fl^ and miR-15/16^fl/fl^ *Cd4*-Cre recipient mice. **I:** Expression by median fluorescent intensity (MFI) using flow cytometry of indicated proteins in spleen Tregs of miR-15/16^fl/fl^ and miR-15/16^fl/fl^ *Cd4*-Cre recipient mice. J: Ratio of total number of FOXP3^+^ Tregs over total number of CD44^hi^CD62L^lo^ T effector cells (Teff) in mesenteric lymph nodes and spleen of miR-15/16^fl/fl^ and miR-15/16^fl/fl^ *Cd4*-Cre Treg recipients. Data from 2 independent experiments. N=5-6 mice/group. Ordinary ANOVA with Dunnett’s multiple comparison test in C-F. 2-way ANOVA with Bonferroni’s multiple comparison test in H, and unpaired t-test 2-tailed in I-J. Bar graphs are shown with error bars demonstrating standard deviation.

### Accumulation of FOXP3^lo^CD25^lo^CD127^hi^ Tregs happens by cell-intrinsic mechanisms

Heightened expansion of miR-15/16 deficient Tregs in the colitis model indicated that the external environment of the host was not driving Treg accumulation and generation of an abnormal Treg phenotype. To study cell intrinsic versus cell extrinsic effects with greater detail, we set up co-cultures of naïve miR-15/16^fl/fl^ and miR-15/16^fl/fl^ *Cd4*-Cre T cells (CD4^+^CD44^lo^CD62L^hi^, distinguishable by the congenic marker CD45.1) that were seeded under Treg-polarizing conditions (Figure 4A). The same number of FOXP3^+^ induced (iTregs) of either genotype accumulated after 5 days in culture (Figure 4B). However, comparisons of co-cultured iTregs showed consistent reduction of FOXP3, CD25 and CTLA4 expression in miR-15/16^fl/fl^ *Cd4*-Cre iTregs compared to miR-15/16^fl/fl^ controls, and CD127 and PD-1 expression were consistently upregulated in miR-15/16 deficient iTregs (Figure 4C). These results confirmed that intrinsic signals drive the observed phenotypic changes in vitro.

**Figure 4.**
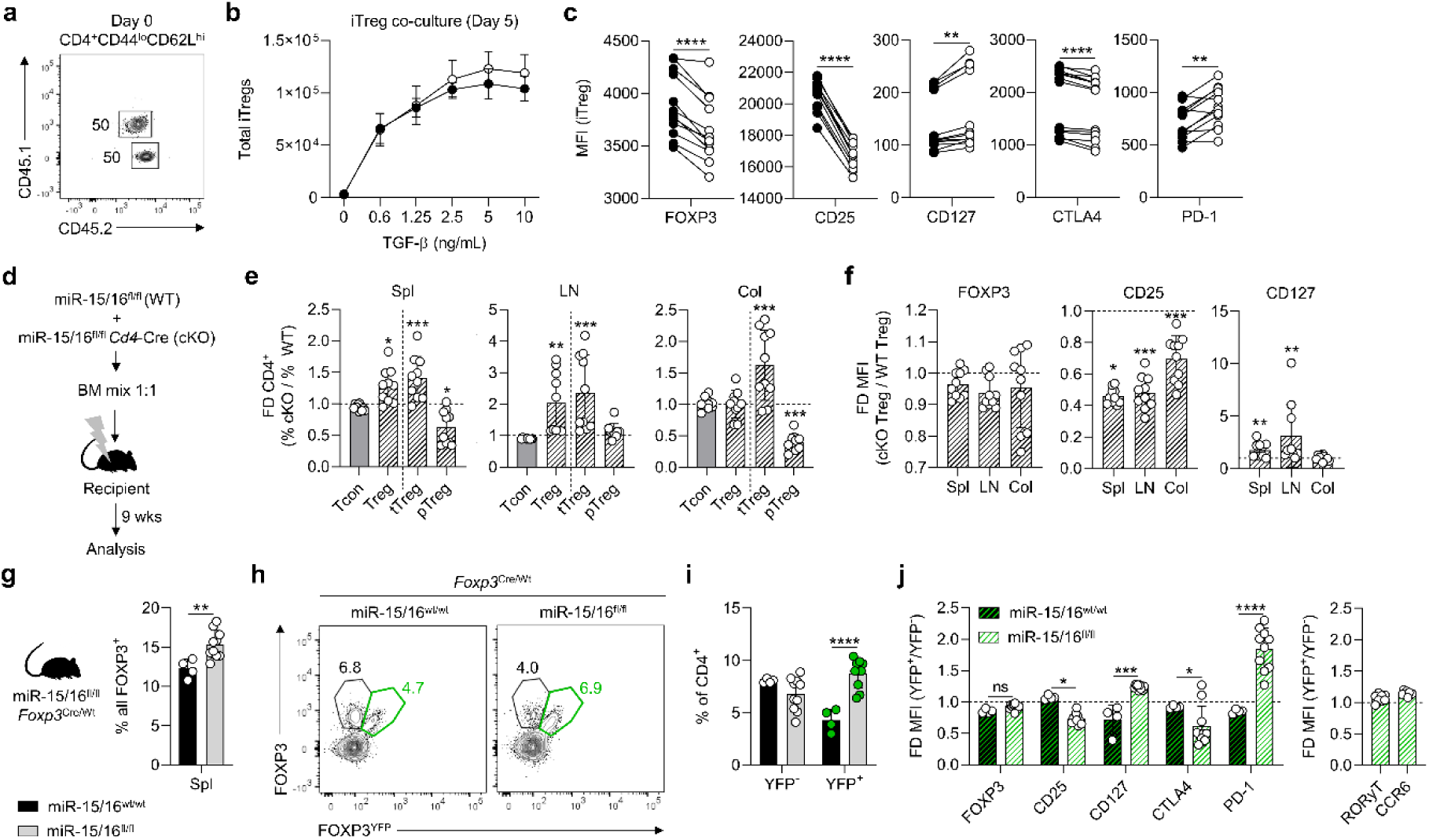
Accumulation of FOXP3^lo^CD25^lo^CD127^hi^ Tregs happens in a cell-intrinsic manner. **A:** Co-cultures of naïve CD4^+^ T cells from CD45 congenic mice (miR-15/16^fl/fl^ and miR-15/16^fl/fl^ *Cd4*-Cre). **B:** Total number of induced Tregs (iTregs) after 5 days in culture under Treg polarizing conditions assessed by FOXP3 expression by flow cytometry. **C:** Protein expression by median fluorescent intensity (MFI); paired analysis of co-cultured iTregs. **D:** Generation of mice with chimeric bone marrow. Recipient mice were analyzed after 9 weeks. Fold difference of cell abundance **(E)** and MFI by flow cytometry of indicated proteins in Tregs **(F)** from three tissues (change in miR-15/16^fl/fl^ *Cd4*-Cre from miR-15/16^fl/fl^ WT control). **G:** Frequency of Tregs among all CD4^+^ T cells in natural chimeric female miR-15/16^fl/fl^ *Foxp3*^Cre/Wt^ mice. **H:** Representative contour plots of gating strategy to separate FOXP3^+^ cells based on YFP reporter signal (indicating Cre expression). YFP^-^ and YFP^+^ Treg frequency among all CD4^+^ T cells **(I)** and fold difference of flow cytometry MFI of indicated proteins expressed in Tregs (change in YFP^+^ Tregs from YFP^-^ Tregs) **(J)**. Data from a minimum of 2 independent experiments. N=4-10 mice/group. Paired t-test 2-tailed in C. Ordinary ANOVA with Dunnett’s multiple comparison test in E-F. 2-way ANOVA with Sidiak’s multiple comparison test in I-J. Bar graphs are shown with error bars demonstrating standard deviation.

We came to the same conclusion in vivo using mixed hematopoietic chimeras. We transferred an equal ratio of miR-15/16^fl/fl^ and miR-15/16^fl/fl^ *Cd4*-Cre bone marrow cells to lethally irradiated recipient mice, and analyzed them after 9 weeks of reconstitution (Figure 4D). miR-15/16^fl/fl^ *Cd4*-Cre Tregs were more abundant than miR-15/16^fl/fl^ control Tregs in spleen and lymph nodes (Figure 4E). Specifically, Helios^+^ tTregs were responsible for expanding the Treg pool. FOXP3^−^ Tcons were found at a 1:1 ratio in all tissues (Figure 4E). FOXP3 expression trended lower in miR-15/16^fl/fl^ *Cd4*-Cre Tregs compared to miR-15/16^fl/fl^ Tregs from the same mice (Figure 4F, left panel), and the expression of CD25 was consistently and robustly reduced in miR-15/16 deficient Tregs (Figure 4F, middle panel). Conversely, CD127 expression was upregulated several fold in spleen and lymph node Tregs lacking miR-15/16 (Figure 4F, right panel).

Finally, we used miR-15/16^fl/fl^*Foxp3*^Cre/Wt^ *Foxp3*-Cre-YFP heterozygous females as a third experimental approach that allowed us to study Cre-expressing miR-15/16 deficient Tregs (YFP^+^) and miR-15/16 sufficient Tregs (YFP^−^) in the same animal without intervention with irradiation and bone marrow transplantation. Despite allelic expression of Cre and YFP due to X-chromosome inactivation, having only one *Foxp3*^Cre^ allele was enough to increase the total number of Tregs in female miR-15/16^fl/fl^*Foxp3*^Cre/Wt^ mice compared to miR-15/16^wt/wt^*Foxp3*^Cre/Wt^ controls (Figure 4G). Separating Tregs by YFP signal showed that these increases reflected a specific increase in the miR-15/16-deficient YFP^+^ Tregs in miR-15/16^fl/fl^*Foxp3*^Cre/Wt^ mice (Figure 4H-I), thereby confirming that miR-15/16 prevent Treg overgrowth in a cell-intrinsic manner. Moreover, these expanded miR-15/16-deficient Tregs also displayed the characteristic low expression of FOXP3, CD25 and CTLA4, and high expression of CD127 and PD-1 (Figure 4J). RORγT and CCR6 expression were similar in miR-15/16 deficient and sufficient Tregs (Figure 4J). Together, these results demonstrate that miR-15/16 regulate gene expression networks in Tregs that limit their proliferation and modulate the expression of hallmark proteins.

### Access to IL-2 does not restore Treg CD25 expression in absence of miR-15/16

IL-2 signaling plays a fundamental role in Treg homeostasis and function, and exposure to IL-2 induces expression of its own high affinity receptor chain, CD25 (a.k.a. IL2Rα) (6,30). The other subunits of the IL2R complex (CD122/IL2Rβ and CD132/IL2Rγ; a.k.a. “common γ chain”) were expressed similarly in Tregs from miR-15/16^fl/fl^ *Cd4*-Cre and miR-15/16^fl/fl^ control mice (Figure 5A-B). To test whether reduced IL-2 availability could be responsible for impaired CD25 expression in miR-15/16 deficient Tregs, we treated co-cultured iTregs from miR-15/16^fl/fl^ and miR-15/16^fl/fl^ *Cd4*-Cre mice with different IL-2 concentrations including IL-2 neutralization by addition of an anti-IL-2 antibody. Neutralizing IL-2 reduced iTreg differentiation, affecting both genotypes similarly (Figure 5C). However, the amount of exogenously provided IL-2 did not alter the number of iTregs nor did it restore CD25 expression in miR-15/16 deficient iTregs (Figure 5C). In fact, miR-15/16 deficient iTregs responded to IL-2 stimulation by upregulating CD25 to the same degree as miR-15/16 sufficient iTregs (Figure 5C), although, starting at a lower expression level prevented them from reaching full CD25 expression. In vivo generated Tregs sorted by FACS and cultured with or without IL-2 overnight confirmed this finding. CD25 expression increased in the presence of IL-2 in both control and miR-15/16 deficient Tregs (Figure 5D), with miR-15/16 deficient Tregs consistently exhibiting lower CD25 expression in all conditions (Figure 5D). These findings demonstrate that miR-15/16 are necessary to secure strong CD25 expression in Tregs independently of access to IL-2.

**Figure 5.**
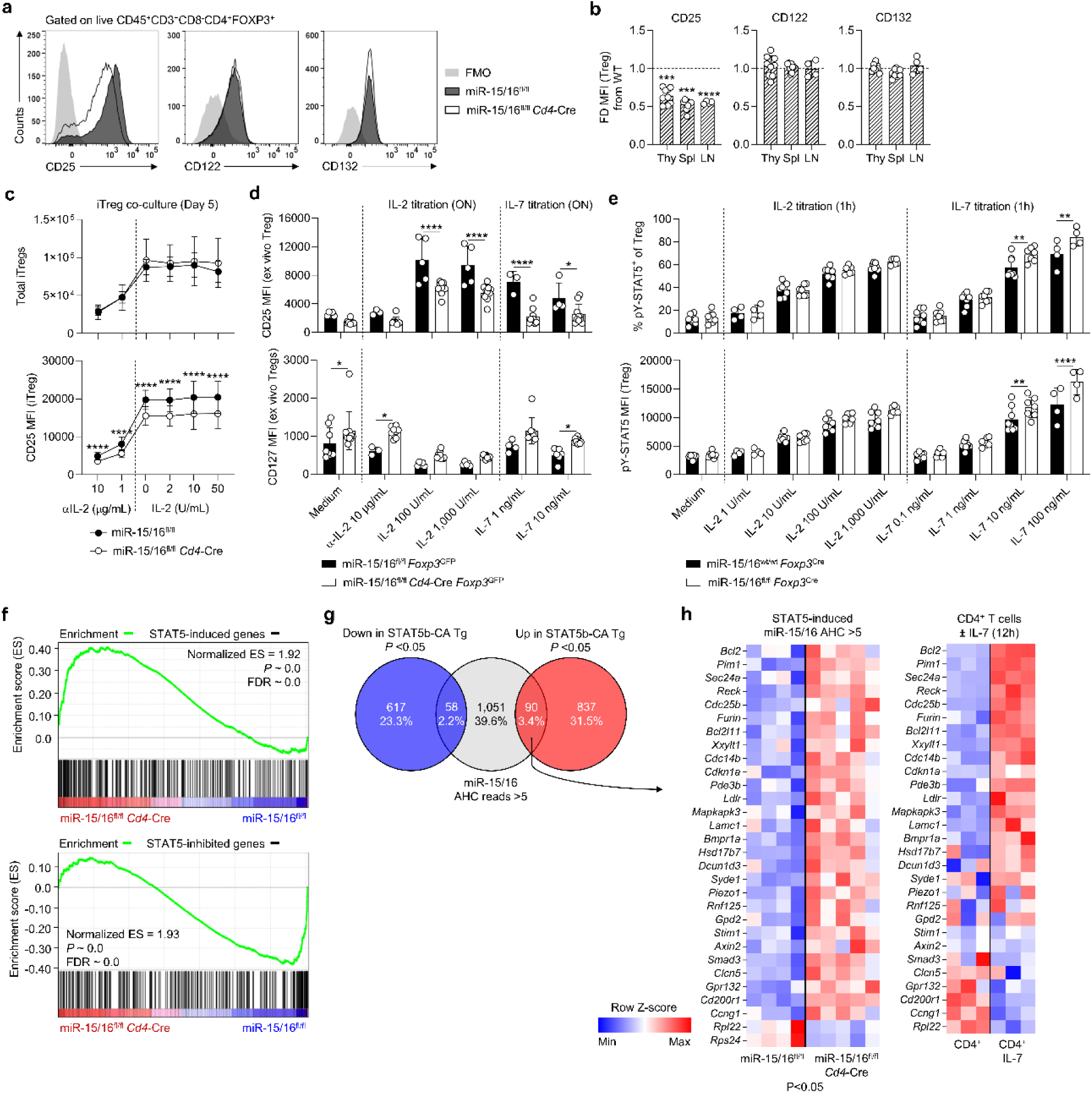
Derepression of miR-15/16 target CD127 promotes activation of STAT5 via IL-7. IL-2R subunit expression in representative flow cytometry histograms **(A)** and fold difference of expression by median fluorescent intensity (MFI) **(B)** in miR-15/16^fl/fl^ and miR-15/16^fl/fl^ *Cd4*-Cre Tregs from indicated tissues (change in miR-15/16^fl/fl^ *Cd4*-Cre from miR-15/16^fl/fl^ WT control). **C:** Total number of cells and CD25 expression by MFI in co-cultures of induced Tregs (iTregs) from CD45 congenic mice (miR-15/16^fl/fl^ and miR-15/16^fl/fl^ *Cd4*-Cre) under different IL-2 concentrations. **D:** CD25 and CD127 expression by MFI in Tregs isolated by FACS and cultured overnight (ON) with IL-2, IL-7 or in medium. **E:** Ex vivo CD4^+^ T cell cultures from miR-15/16^wt/wt^ *Foxp3*^Cre^ and miR-15/16^fl/fl^ *Foxp3*^Cre^ mice stimulated 1h with IL-2, IL-7 or unstimulated medium control followed by flow cytometry analysis of frequency of Tregs with phosphorylated STAT5 (anti-pY694-STAT5) and Treg pSTAT5 MFI. **F:** Gene set enrichment analysis (GSEA) of genes upregulated (‘STAT5-induced genes’) or downregulated (‘STAT5-inhibited genes’) in Tregs with constitutively active STAT5b (extracted from (6)) in CD4^+^ T cells from miR-15/16^fl/fl^ *Cd4*-Cre mice and miR-15/16^fl/fl^ control mice (22). **G:** Venn diagram demonstrating overlap between STAT5-inhibited genes (‘Down in STAT5b-CA Tg’), STAT5-induced genes (‘Up in STAT5b-CA Tg’) and putative miR-15/16 target genes identified by Ago2 high-throughput sequencing of RNAs isolated by crosslinking immunoprecipitation (AHC) with read depth >5 (22). **H:** Expression of overlap genes (‘miR-15/16 AHC reads >5’ and ‘Up in STAT5b-CA Tg’) in CD4^+^ T cells of miR-15/16^fl/fl^ and miR-15/16^fl/fl^ *Cd4*-Cre mice by RNA-seq (22), and their expression in CD4^+^ T cells after IL-7 stimulation (52). Data from a minimum of 2 independent experiments. N=4-9/group. Ordinary ANOVA with Dunnett’s multiple comparison test in B. 2-way ANOVA with Sidiak’s multiple comparison test in C-E. Bar graphs are shown with error bars demonstrating standard deviation.

### Derepression of the miR-15/16 target CD127 promotes IL-7-dependent STAT5 activation

Tregs and Tcons display reciprocal expression of CD25 and CD127, with Tregs presenting as CD25^hi^CD127^lo^ and Tcons as CD25^lo^CD127^hi^. Consequently, IL-2 and IL-7 have opposing limiting functions in T cell immunity, with IL-2 required to maintain tolerance and IL-7 important for supporting immunological memory (31). Since miR-15/16 was required to maintain the CD25/CD127 balance in Tregs, we studied the effect of IL-7 in our ex vivo Treg cultures. IL-7 promoted T cell survival regardless of genotype (data not shown), with little direct effect on CD25 and CD127 expression (Figure 5D). Next, we treated cells with IL-2 or IL-7 ex vivo and measured STAT5 phosphorylation (pSTAT5) in responding cells by flow cytometry. Despite the fact that miR-15/16 expression is required for high CD25 expression in Tregs, the frequency of pSTAT5^+^ cells and pSTAT5 signal intensity (MFI) were similar in miR-15/16 deficient and control Tregs in response to IL-2 stimulation (Figure 5E). In contrast, IL-7 stimulation selectively increased the frequency of pSTAT5^+^ cells and pSTAT5 MFI in miR-15/16 deficient Tregs, especially at high cytokine concentrations (Figure 5E).

IL-2R deficient Treg function can be rescued by constitutive STAT5 expression (6). We extracted a list of differentially expressed genes by RNA-seq of Tregs from transgenic STAT5b-CA mice and control Tregs and analyzed STAT5-induced or STAT5-inhibited genes in CD4^+^ T cells isolated from miR-15/16^fl/fl^ *Cd4*-Cre and miR-15/16^fl/fl^ control mice. We identified a significant enrichment of STAT5-induced genes in miR-15/16^fl/fl^ *Cd4*-Cre T cells, and an enrichment of STAT5-inhibited genes in miR-15/16^fl/fl^ control T cells (Figure 5F). Genes directly targeted by miR-15/16 in T cells were analyzed by Ago2 high-throughput sequencing of RNAs isolated by crosslinking immunoprecipitation (AHC) in our previous study (22). Putative miR-15/16 targets were identified as genes with one or more miR-15/16 3’UTR seed-matches corresponding with AHC read depth >5, and this gene set was highly enriched among genes upregulated in miR-15/16^fl/fl^ *Cd4*-Cre T cells (22). Interestingly, this set of putative miR-15/16 targets significantly overlapped with genes upregulated in Tregs with constitutive STAT5 expression (Figure 5G). Furthermore, several of these STAT5-induced putative miR-15/16 target genes were expressed at a significantly higher level in miR-15/16 deficient T cells compared to miR-15/16^fl/fl^ controls (Figure 5H), and a majority were induced in response to IL-7 stimulation (Figure 5G). Collectively, these findings suggest that IL-7 promotes STAT5-dependent gene expression programs that are targeted by miR-15/16 in Tregs, and that miR-15/16 regulate Treg homeostasis by coordinating cytokine responsiveness and cytokine-induced gene expression programs.

### Treg overgrowth can compensate for impaired suppressive capacity

The inflammatory reactions happening systemically in miR-15/16^fl/fl^ *Foxp3*^Cre^ mice indicate that Treg suppression is compromised in the absence miR-15/16, despite the increased number of Tregs in these mice. To measure the suppressive capacity of miR-15/16-deficient Tregs, we used in vitro suppression assays. CD4^+^CD25^-^CD45.1^+^ responder Tcons were sorted from wildtype mice and activated in presence of CD4^+^GFP^+^CD45.1^-^ Tregs sorted from miR-15/16^fl/fl^ *Cd4*-Cre *Foxp3*^GFP^ and miR-15/16^fl/fl^ *Foxp3*^GFP^ control mice. Surprisingly, Tcons cultured with miR-15/16^fl/fl^ *Cd4*-Cre Tregs or miR-15/16^fl/fl^ control Tregs proliferated similarly, as indicated by CellTrace^TM^ Violet (CTV) dye dilution (Figure 6A-B), suggesting that Tcons experienced the same degree of suppression by Tregs of either genotype. However, we analyzed Treg frequency 72h after seeding the cells and found a significant increase in the number of miR-15/16^fl/fl^ *Cd4*-Cre Tregs compared to control Tregs which was enhanced in wells with responder cells present (Figure 6C). Labeling the Tregs with CTV confirmed increased proliferation in the absence of miR-15/16 (Figure 6D-E). We therefore recalculated the suppression profile, switching from the seeding ratio at the start of the culture to the final Treg:Tcon ratio at 72h ((Figure 6F). This analysis revealed that a greater number of Tregs from miR-15/16^fl/fl^ *Cd4*-Cre mice were needed to achieve the same degree of suppression as Tregs from miR-15/16^fl/fl^ control mice. Using linear regression analysis, we estimate 15% lower suppression by miR-15/16^fl/fl^ *Cd4*-Cre Tregs (Figure 6F). These results demonstrate that miR-15/16 restrict Treg expansion ex vivo, and that excessive expansion due to miR-15/16 deficiency can compensate for lower suppressive capacity on a per-cell-basis.

**Figure 6.**
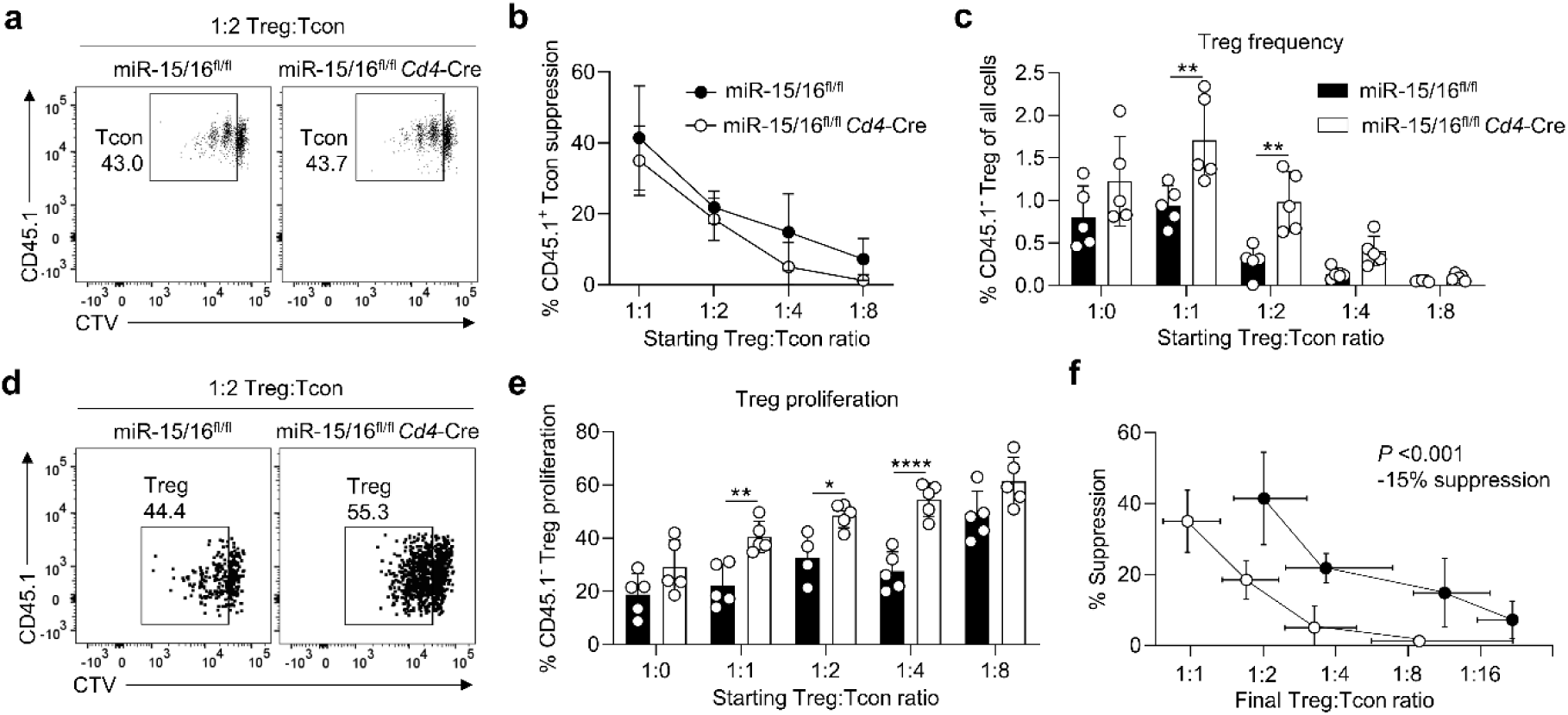
Treg overgrowth, caused by miR-15/16 deficiency, compensates for impaired suppressive ability. **A:** Representative dot plots of CellTrace^TM^ Violet (CTV) dilution in CD4^+^CD25^-^CD45.1^+^ responder cells (Tcons) from WT mice, cultured with CD4^+^FOXP3-GFP^+^CD45.1^-^ Tregs from miR-15/16^fl/fl^ *Cd4*-Cre *Foxp3*^GFP^ mice or miR-15/16^fl/fl^ *Foxp3*^GFP^ control mice. **B:** Quantification of Treg suppression calculated based on Treg:Tcon seeding ratio at start of the culture. **C:** Treg frequency among all T cells. **D:** Representative dot plots of CTV dilution CD4^+^FOXP3-GFP^+^CD45.1^-^ Tregs from miR-15/16^fl/fl^ *Cd4*-Cre *Foxp3*^GFP^ and miR-15/16^fl/fl^ *Foxp3*^GFP^ control. **E:** Quantification of Treg proliferation (i.e. frequency of Tregs dividing ≥1). **F:** Quantification of Treg suppression calculated based on Treg:Tcon seeding ratio at the end of the culture (72h post seeding).

### miR-15/16 deficient Tregs resemble naturally occurring CD25^lo^ Tregs

To learn more about the impact of miR-15/16 regulation on Treg transcriptional programs, we isolated Tregs from miR-15/16^fl/fl^*Foxp3*^Cre^ mice (‘cKO Treg’) and miR-15/16^wt/wt^*Foxp3*^Cre^ controls (‘WT Treg’) for RNA-seq analysis (Figure 7A). From miR-15/16^wt/wt^*Foxp3*^Cre^ mice, we additionally sorted CD25^hi^ and CD25^lo^ Tregs (Figure 7A and supplementary Table S1). 168 genes were differentially expressed in CD25^hi^ versus CD25^lo^ Tregs and in WT versus cKO Tregs (Figure 7B). Differentially expressed genes (DEG) included several known to modulate Treg function (e.g. *Myc*, *Myb*, *Bcl6*, *Tgfbr1*, *Ikzf2* (Helios) and *Ikzf4* (Eos)). Interestingly, the transcriptional profile of cKO Tregs resembled that of naturally occurring CD25^lo^ WT Tregs (Figure 7B). Mapping our AHC data onto the Treg DEGs revealed an enrichment for AHC reads corresponding with miR-15/16 seed sequences within the 3’ UTR of genes upregulated in cKO Tregs, compared with those more highly expressed in WT Tregs (Figure 7B, bars on right side). Quantifying this enrichment showed that significantly more genes upregulated in cKO Tregs had AHC reads ≥5 at miR-15/16 target sites (Figure 7C). Thus, miR-15/16 directly target some, but not all of the DEGs shared by cKO Tregs and WT CD25^lo^ Tregs. A full transcriptome-wide analysis of cKO and WT Tregs demonstrated that miR-15/16 bind and regulate a large number of direct RNA targets in Tregs (Figure S3).

**Figure 7.**
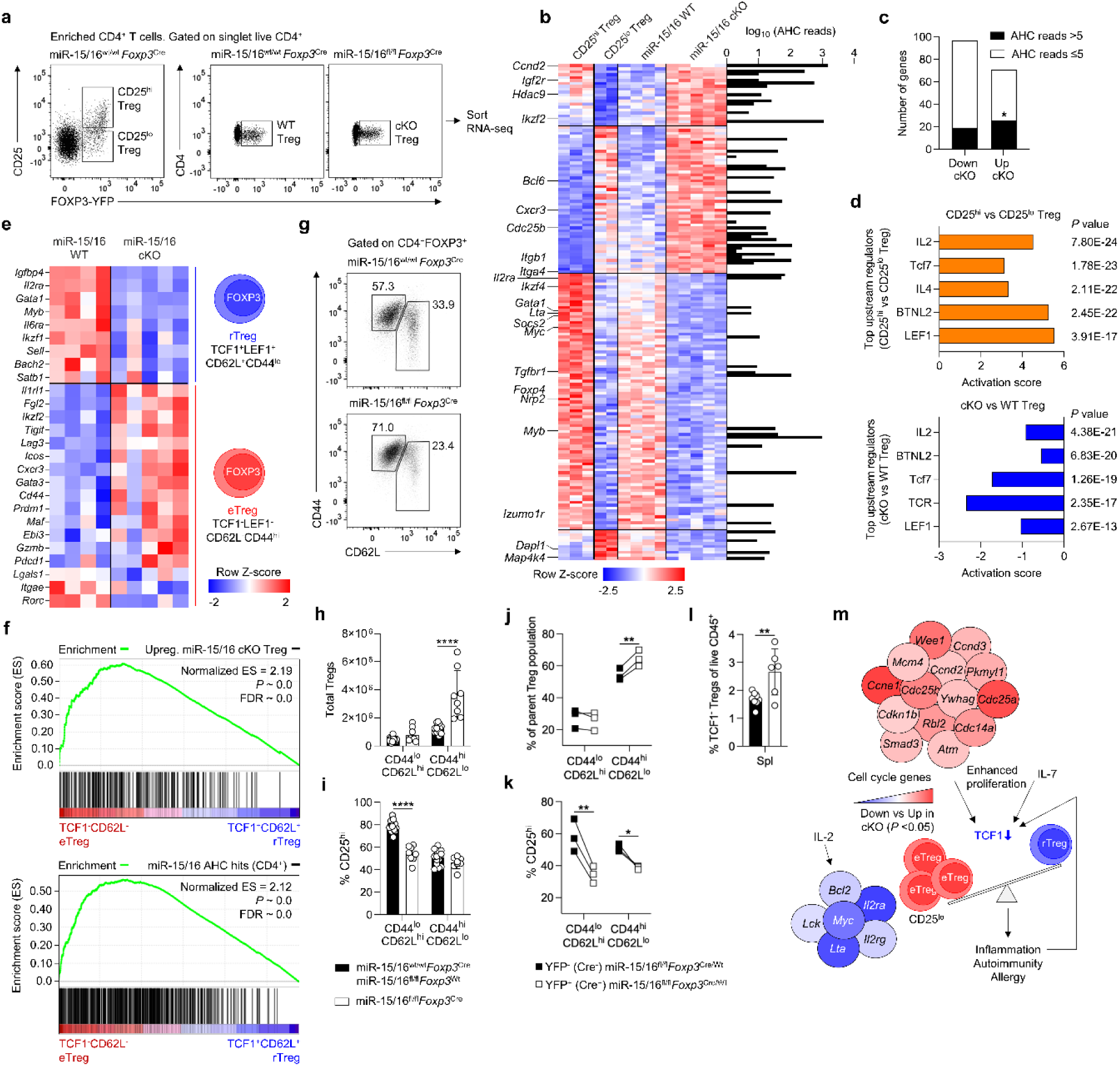
miR-15/16-mediated suppression of cell cycle genes prevents downregulation of TCF1 and expansion of TCF1^-^ effector Tregs. **A:** Flow cytometry dot plots of Treg populations isolated for RNA-seq analysis (CD25^hi^ Tregs, CD25^lo^ Tregs, WT Tregs and miR-15/16 cKO Tregs). **B:** Heatmap of differentially expressed genes (DEG) in CD25^hi^ versus CD25^lo^ Tregs and WT versus cKO Tregs (P value <0.05). The whole DEG list is provided as supplementary Table S1. The heatmap is plotted alongside a bar graph of AHC read depth at miR-15/16 seed-matches for each gene at which they occur (22). **C:** Genes with AHC reads >5 or AHC reads ≤5 at miR-15/16 seed-matches that are downregulated and upregulated in cKO Tregs compared to WT Tregs (22). **D:** Upstream regulators by Ingenuity Pathway Analysis of DEGs in CD25^hi^ Tregs versus CD25^lo^ Tregs (top) and miR-15/16 cKO Tregs versus WT Tregs (bottom). **E:** Heatmap expression of transcription factors and key functional molecules that distinguish resting Treg (rTreg; TCF1^+^LEF1^+^CD62L^+^CD44^lo^) and effector Treg (eTreg; TCF1^-^LEF1^-^CD62L^-^CD44^hi^) subgroups described by (11). Expression in WT Tregs and miR-15/16 cKO Tregs is shown. **F:** Gene set enrichment analysis (GSEA) of genes upregulated in cKO Tregs (top) (compared to WT) and AHC hits (reads >5) at miR-15/16 seed-matches (bottom) across eTreg and rTreg transcriptomes (11,22). Representative dot plots **(G)** and quantification of total **(H)** CD62L^hi^CD44^lo^ Tregs and CD62L^lo^CD44^hi^ Tregs in miR-15/16^fl/fl^ *Foxp3*^Cre^ mice and miR-15/16^wt/wt^ *Foxp3*^Cre^ and miR-15/16^fl/fl^ *Foxp3*^Wt^ control mice. **I:** Frequency of CD25^hi^ cells among CD62L^hi^CD44^lo^ Tregs and CD62L^lo^CD44^hi^ Tregs in the same mice. **J:** Frequency of CD62L^hi^CD44^lo^ and CD62L^lo^CD44^hi^ Tregs among all Tregs in heterozygous female miR-15/16^fl/fl^ *Foxp3*^Cre/Wt^ mice and frequency of CD25 expressing cells among the same Treg populations in the same mice **(K)**. **L:** Frequency of TCF1^-^ Treg among live hematopoietic cells in spleen of miR-15/16^fl/fl^ *Foxp3*^Cre^ mice and miR-15/16^wt/wt^ *Foxp3*^Cre^ and miR-15/16^fl/fl^ *Foxp3*^Wt^ control mice. **M:** Schematic illustration of proposed mechanism where enhanced proliferation due to increased cell cycle gene expression leads to TCF1 downregulation to promote eTreg generation. eTregs, characterized by lower CD25 expression, have reduced IL-2 induced gene expression and insufficiently suppress immune activation. Data from a minimum of 2 independent experiments. N=2-10 mice/group. Chi square test in C. 2-way ANOVA with Sidiak’s multiple comparison test in H-I. Paired t-test 2-tailed in J-L. Bar graphs are shown with error bars demonstrating standard deviation.

### miR-15/16 maintain a resting Treg transcriptional signature

Next, we used Ingenuity Pathway Analysis (IPA) to analyze all DEGs in CD25^hi^ versus CD25^lo^ Tregs to generate a list of candidate upstream regulators (Figure 7D). The same analysis performed on all DEGs in cKO versus WT Tregs identified several of the same upstream regulators, including *Il2*, *Tcf7* (TCF1) and *Lef1* (Figure 7D). Expression of the transcription factor TCF1 and its binding partner LEF1 was recently shown to separate Tregs into distinct subpopulations; activated Tregs, effector Tregs (eTregs) and resting Tregs (rTreg) (11). Therefore, we analyzed the expression of transcription factors and key functional molecules that distinguish TCF1^+^LEF1^+^CD62L^+^CD44^lo^ rTregs from TCF1^-^LEF1^-^ CD62L^-^CD44^hi^ eTregs in miR-15/16^fl/fl^ *Foxp3*^Cre^ and miR-15/16^wt/wt^ *Foxp3*^Cre^ mice. The transcriptional profile of rTregs resembled that of WT Tregs, whereas eTregs corresponded more closely with cKO Tregs (Figure 7E). Expanding analysis to the entire transcriptome of rTregs and eTregs showed that genes upregulated in miR-15/16 cKO Tregs (Figure 7F, top) and miR-15/16 AHC hits (Figure 7F, bottom) were enriched in the eTreg transcriptional profile. Together, these results suggest that miR-15/16 maintain distinct subsets of Tregs with a resting phenotype.

### miR-15/16 restrict the formation of eTregs

In line with the eTreg transcriptional signature of miR-15/16 cKO Tregs, CD44^hi^CD62L^lo^ Tregs were selectively expanded in miR-15/16^fl/fl^ *Foxp3*^Cre^ mice compared to miR-15/16^wt/wt^ *Foxp3*^Cre^ controls (Figure 7G-H). CD25 expression was markedly reduced among CD44^lo^CD62L^hi^ Tregs lacking miR-15/16, mimicking the lower frequency of CD25^hi^ cells in the CD44^hi^CD62L^lo^ Treg subset (Figure 7I). Analysis of the same Treg subsets in miR-15/16^fl/fl^*Foxp3*^Cre/Wt^ *Foxp3*-Cre-YFP heterozygous female mice confirmed that miR-15/16 deficiency promotes a selective increase of CD44^hi^CD62L^lo^ Tregs (Figure 7J) with reduced CD25 expression most pronounced in the CD44^lo^CD62L^hi^ Treg subset (Figure 7K). TCF1 protein expression was reduced in the whole Treg pool of miR-15/16^fl/fl^ *Foxp3*^Cre^ mice, supporting a shift in transcriptional programs leading to expansion of TCF1^-^ eTregs (Figure 7L).

In summary, miR-15/16 deficient Tregs exhibited an effector Treg transcriptional signature and an increased proportion of cells with concordant expression of eTreg marker proteins. We propose a model wherein increased cell cycle and IL-7 signaling in the absence of miR-15/16 downregulates TCF1 expression that, in turn, promotes formation of eTregs that are characterized by lower CD25 expression, and hence decreased expression of IL-2-induced genes (Figure 7M). eTreg overgrowth reduces the diversity of specialized suppressive Treg functions to an extent that tissue homeostasis cannot be preserved and leads to immune activation and inflammation.

## Discussion

Several decades of research have established Tregs as key suppressors of the immune system (32), where FOXP3 and IL-2 signaling via STAT5 orchestrate the core events that define and reinforce Treg identity (6). In more recent years, Treg heterogeneity has been explored (11,33,34), demonstrating that specific transcription factors, TCR repertoires and cytokine signals drive functional specialization of Treg subsets (11,35,36). miRNAs exert their biologic effects through multiple targets in gene networks which makes them effective in regulating cellular trajectories (22,37,38). The experiments in this study revealed that miR-15/16 play a critical role in the regulation of Treg behavior. miR-15/16 expression was essential to develop a normal Treg phenotype under homeostatic conditions, to limit their proliferative capacity and efficiently suppress CD4^+^ Teff cells. Treg-specific ablation of miR-15/16 resulted in non-fatal but substantial systemic inflammatory responses in mice.

It remains unclear why Tregs fail to suppress the immune system in chronic inflammatory diseases, and there is a need to increase our knowledge of the cellular programs capable of limiting inflammation and restoring tissue homeostasis. Experiments using OVA challenge of the airways to provoke type 2 inflammation elicited significantly more eosinophils in miR-15/16^fl/fl^ *Foxp3*^Cre^ mice compared to WT controls, despite miR-15/16^fl/fl^ *Foxp3*^Cre^ mice having a higher number of airway-resident Tregs. This disconnect between Tregs numbers and physiological function highlights the importance of functional specialization in Treg suppression of immune activation. Spontaneous tissue inflammation prior to allergen challenge may have affected the ability of Tregs to suppress additional inflammatory cues induced by OVA in miR-15/16^fl/fl^ *Foxp3*^Cre^ mice. Treg exhaustion has frequently been reported in malignancies where checkpoint blockade through PD-1 is an effective tumor treatment (39). miR-15/16^fl/fl^ *Foxp3*^Cre^ Tregs were characterized by high PD-1 expression, a feature associated with exhaustion, IFN-γ production and reduced suppression of CD4^+^ Teff cells (40). miR-15/16^fl/fl^ *Foxp3*^Cre^ mice displayed increased numbers of Teff cells and elevated IFN-γ production, indicating possible Treg dysfunction related to cell exhaustion.

miR-15/16 are known tumor suppressors, best characterized in B cell leukemia, that affect cell growth by restricting cell cycle and anti-apoptotic genes such as *Ccnd1* (Cyclin D1), *Bmi1, Bcl2*, and *Mcl1* (21,41). Our previous work demonstrated that miR-15/16 also regulate CD8^+^ T cell expansion and differentiation through a large network of genes that control cell cycle, memory and survival (22). The current study revealed a discrepancy in the CD4^+^ T helper cell compartment, where miR-15/16 restrict expansion of Tregs but not Tcons. Using available human data, we identified differences in the abundance of miR-15/16 family miRNAs in defined CD4^+^ T helper cell subsets, with a striking high expression in Tregs compared to conventional Th1, Th2, Th17 and naïve T cells. This observation may indicate more substantial dependence on miR-15/16 regulation in Tregs and is supported by a report of enhanced pTreg generation from miR-15/16-overexpressing conventional CD4^+^ T cells in vivo (27). Furthermore, we showed that miR-15/16 modulated expression of key Treg functional proteins. Throughout the study, miR-15/16 deficient Tregs were characterized as FOXP3^lo^CD25^lo^CD127^hi^ with lower CTLA4, and higher PD-1 expression compared to miR-15/16 sufficient cells, a phenotype established by cell-intrinsic regulation. More subtle changes in protein expression were observed in Tcons obtained from miR-15/16^fl/fl^ *Cd4*-Cre mice, however, secondary effects on the Tcon phenotype cannot be ruled out since Treg function was altered in those mice. Nevertheless, expression of the direct miR-15/16 target CD127 was altered in Tcons. While CD127 increased dramatically in miR-15/16-deficient Tregs, a subtle increase was observed in Tcons, possibly reflecting lower miR-15/16 activity in the latter. CD127 stimulation, via IL-7 or TSLP, is essential for Treg development (42), and IL-7 stimulation has been shown to temporarily rescue survival of Tregs after inducible CD25 ablation in vivo (43). In our study, CD127 upregulation in miR-15/16 deficient Tregs may have contributed to enhanced survival, while compensating for loss in IL-2-dependent survival due to impaired CD25 expression.

miR-15/16 regulation of Treg expansion was selective to tTregs, not pTregs, despite the subsets sharing a functional phenotype (FOXP3^lo^CD25^lo^CD127^hi^CTLA4^lo^PD-1^hi^). Similarly, in vitro generated iTregs did not exhibit excessive expansion in absence of miR-15/16 as compared to co-cultured miR-15/16 sufficient iTreg controls, confirming selective miR-15/16 control of tTreg expansion. The molecular determinants that govern Treg development in the thymus versus the periphery are currently not clear. However, a study aimed at addressing these questions recently reported that CD25-signaling was essential for early expansion of tTregs and Treg homeostasis in the periphery, whereas lineage stability of mature Tregs was largely CD25-independent (7). As expected, transcriptional analysis of CD25 cKO tTregs (*Foxp3*^Cre^) revealed substantial changes in gene expression programs compared to WT Tregs, including reduced levels of most Treg functional molecules. Interestingly, they observed nearly normal levels of FOXP3 protein in tTregs which suggests that STAT5-regulation of FOXP3 expression in the thymus was largely IL-2-indepenent (7). This result is consistent with previous reports of redundant STAT5 phosphorylation in developing Tregs by other STAT5-activating cytokines such as IL-7 and IL-15 (42,44,45). In our study, enhanced Treg expression of CD127 may have promoted early Treg development which contributed to expansion of the tTreg pool. Indeed, IL-7 stimulation resulted in increased pSTAT5^+^ cell in miR-15/16 deficient CD127^hi^ Tregs and we demonstrated that IL-7-responsive STAT5-induced genes overlapped with miR-15/16-regulated gene sets in CD4^+^ T cells, which included empirically tested direct miR-15/16 targets by AHC (22). In fact, STAT5a binds near the miR-15b/16-2 cluster, inducing transcription of STAT5-target genes while suppressing miR-15b/16-2 expression (46), suggesting that miR-15/16 may participate in a feed-forward circuit that enhances STAT5 transcriptional responses.

One of the most striking observations in our study is that miR-15/16 secure high CD25 expression in Tregs. IL-2 stimulation of ex vivo cultured Tregs demonstrated overall impaired CD25 in the absence of miR-15/16, but confirmed a functionally responsive *Il2ra* locus with expression dynamics similar to those of WT Tregs. CD25 function was further confirmed by normal generation of pSTAT5^+^ cells upon IL-2 stimulation. Nevertheless, the transcription profile of miR-15/16-deficient Tregs resembled that of naturally occurring CD25^lo^ Tregs in WT mice. CD25 expression in Tregs declines progressively with age and is associated with development of autoimmune diseases and immune senescence (43,47,48). Thus, Treg-specific miR-15/16 deficiency may be a useful model for age-related inflammation.

In our transcriptional analysis, the striking overlap of upstream regulators in miR-15/16 deficient Tregs and CD25^lo^ Tregs from WT mice revealed a strong association of TCF1 with miR-15/16 gene networks and CD25 signaling. Other reports also suggest close integration of these pathways, for example, cell cycle progression is regulated by miR-15/16, upregulated in CD25-deficient Tregs (7), and promotes generation of TCF1^-^cells (49). In addition, IL-7 signaling inhibits TCF1 expression in thymocytes and mature T cells (50), and TCF1 deficiency in mice leads to increased thymic output of Tregs (51). While TCF1^-^LEF1^-^ Tregs exhibit active cell cycle and increased proliferation, their suppressive capacity has been reported to remain intact (11). In this study, we observed situations where miR-15/16 exerted compensatory effects on Treg control of CD4^+^ T cell responses. For instance, miR-15/16-deficient Tregs proved capable of suppressing Tcon proliferation in vitro, but closer inspection revealed that increased Treg proliferation compensated for lower suppressive capacity on a per-cell-basis. The same situation appeared in the in vivo cell transfer system, where miR-15/16-deficient Tregs protected mice from intestinal inflammation, while also expanding to greater numbers compared with WT Tregs. Furthermore, miR-15/16 depletion in all T cells did not result in inflammation and autoimmune disease in *Cd4*-Cre mice, but conditional depletion of miR-15/16 in Tregs did. The onset of autoimmune disease in *Foxp3*^Cre^ mice suggests that the shift in the Treg pool to a higher proportion eTregs was not sufficient to keep autoimmunity in check, and our data supports previous reports that demonstrate the importance of having functionally diverse Treg subsets to prevent disease (11).

We conclude that miR-15/16 represents a critical node in the coordinate regulation of the heterogeneity, expansion and suppressive function of the Treg pool. Reducing miR-15/16 expression might be beneficial in cancer immunotherapies where decreased IL-2 sensitivity in Tregs could limit IL-2-induced Treg amplification while maintaining IL-2-responsiveness in tumor infiltrating lymphocytes. Further investigation of miR-15/16 regulation of Tregs may uncover therapeutic targets to modulate Treg suppression in the setting of cancer or autoimmunity.

## Methods

Key resources are listed below.

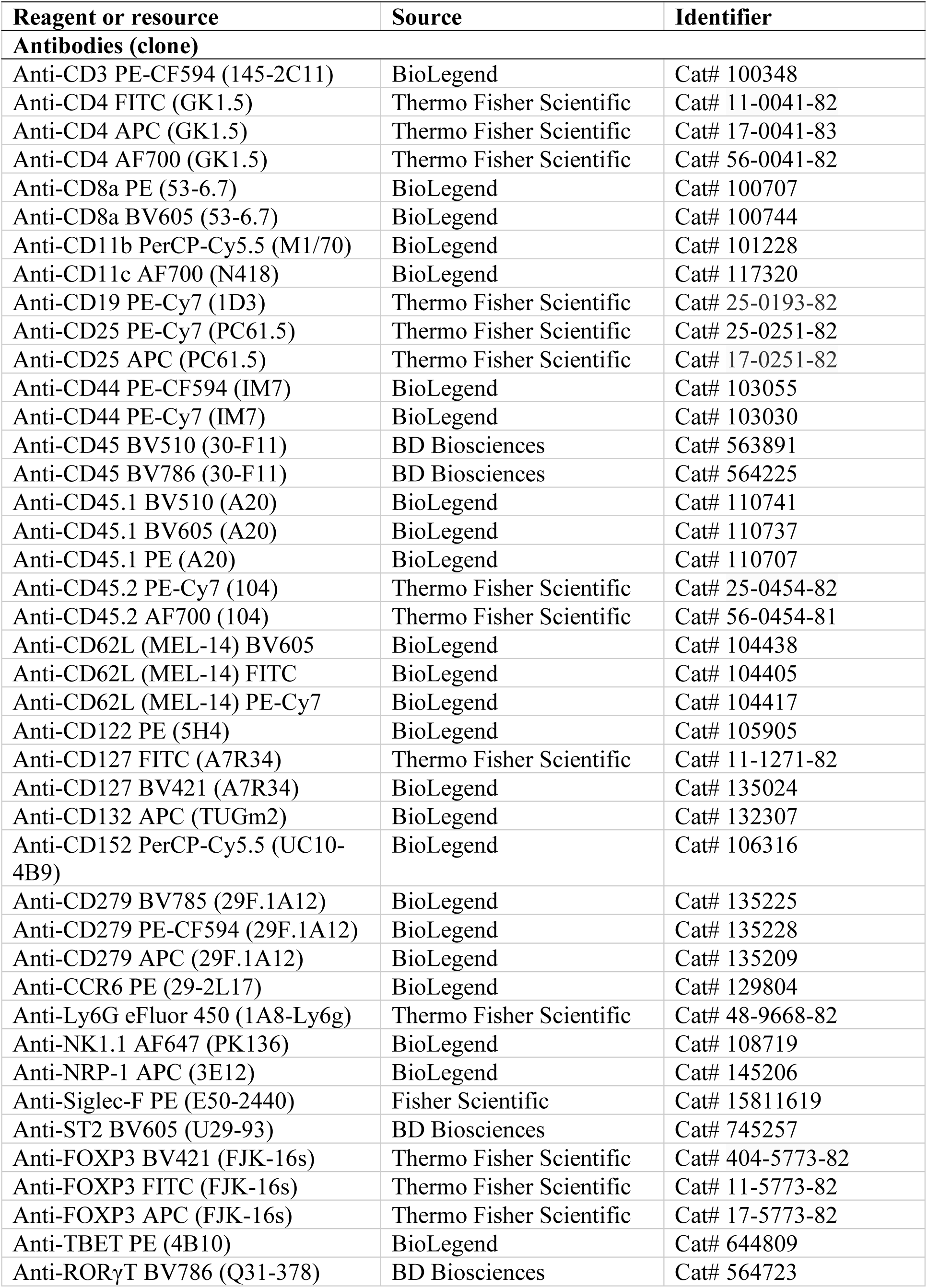
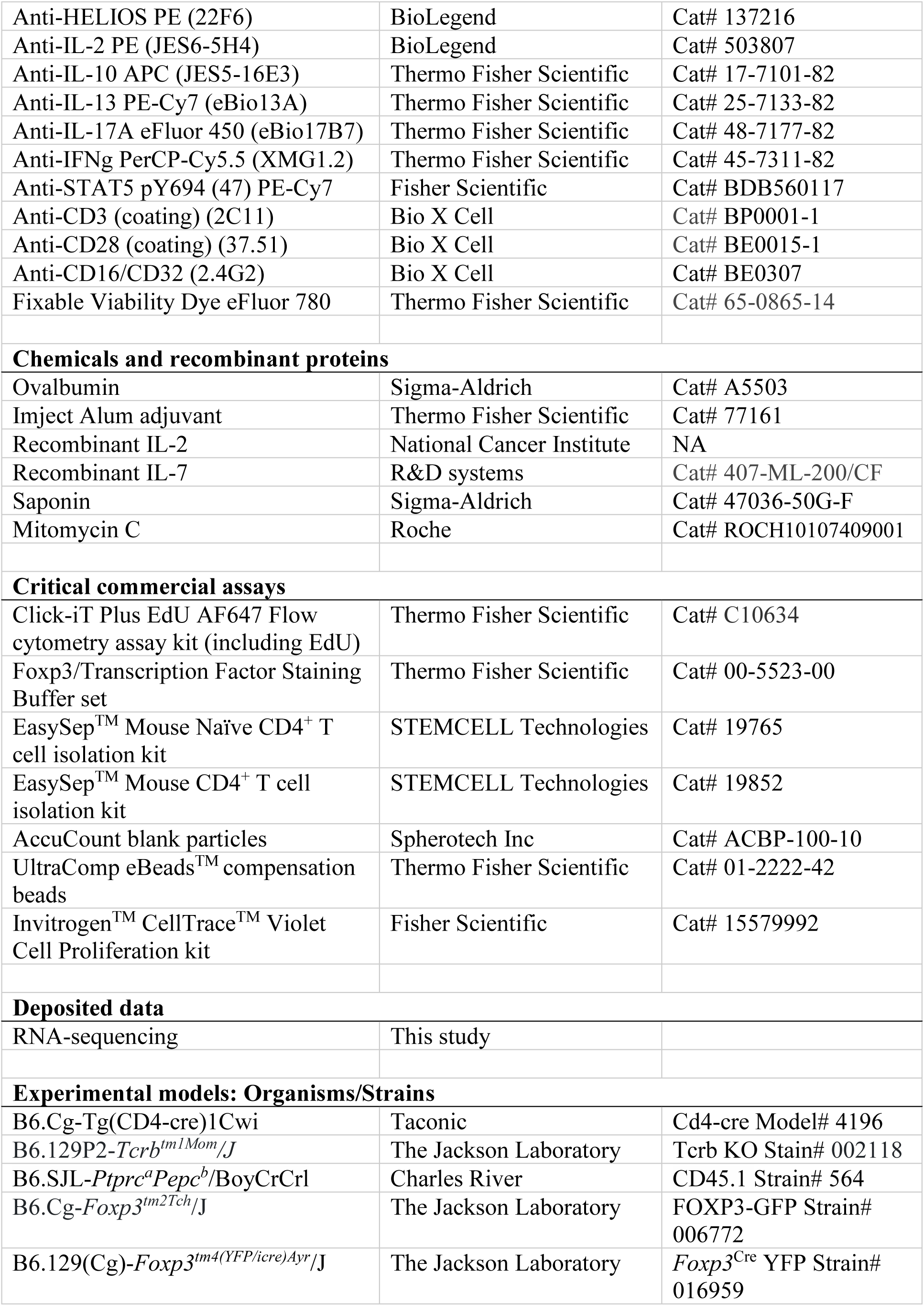
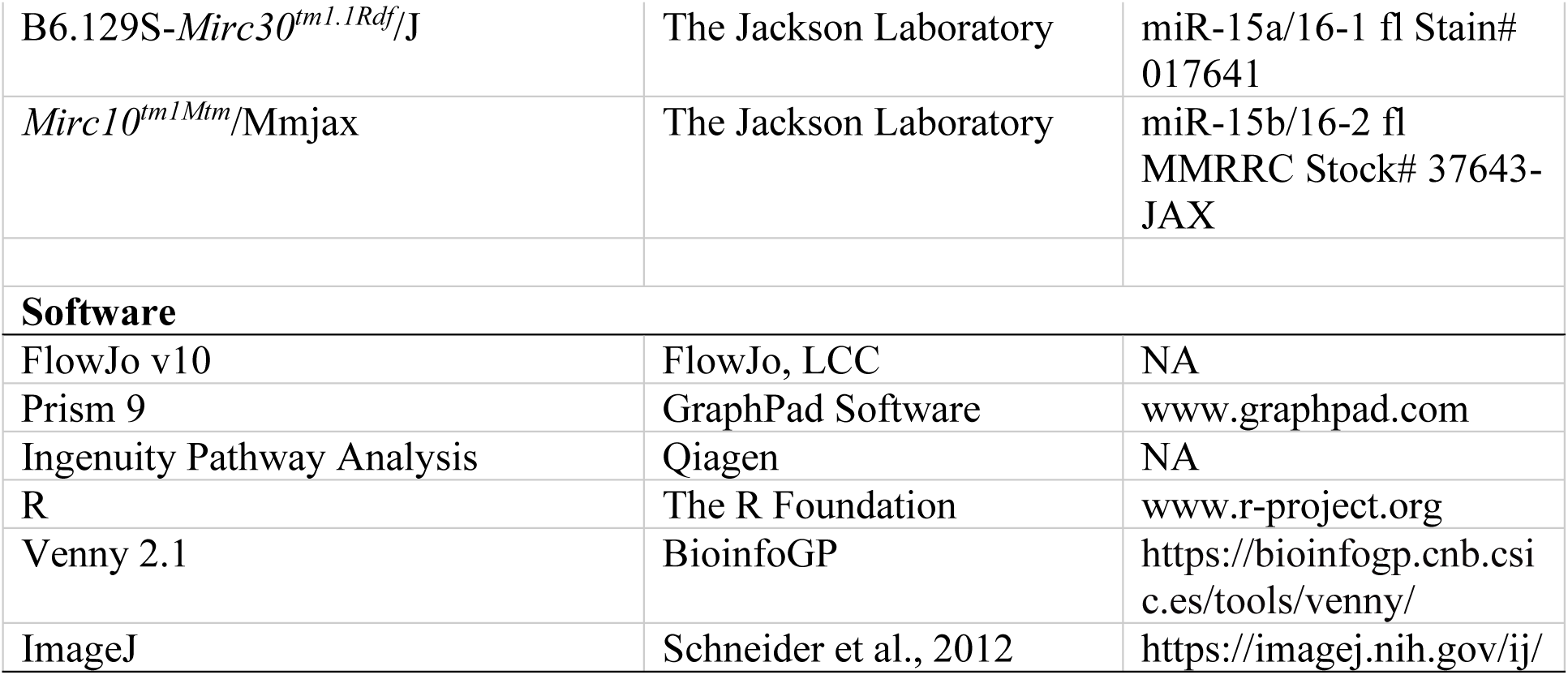

### Lead Contact for reagent and resource sharing

Further information and requests for resources and reagents should be directed to and will be fulfilled by the Lead Contact K. Mark Ansel.

### Mice

miR-15/16^fl/fl^ *Cd4*-Cre mice with conditional inactivation of miR-15/16 family miRNAs (miR-15a, miR-15b, miR-16-1, miR-16-2) in T cells, inactivation of single miR-15a/16-1 or miR-15b/16-2 clusters in T cells, and mice for hematopoietic chimeras were generated as described previously (22). miR-15/16^fl/fl^ *Foxp3*^Cre^ mice with conditional inactivation of miR-15/16 family miRNAs in Tregs were generated by crossing miR-15/16^fl/fl^ mice (loxP-flanked miR-15a/16-1 and miR-15b/16-2 alleles (22)) with B6.129(Cg)-*Foxp3^tm4(YFP/icre)Ayr^*/J mice, kindly provided by the Rosenblum lab (University of California San Francisco). miR-15/16^fl/fl^ *Cd4*-Cre mice carrying a FOXP3 GFP-reporter were generated by crossing miR-15/16^fl/fl^ *Cd4*-Cre mice (22) with B6.Cg-*Foxp3^tm2Tch^*/J mice. In all experiments, male and female age and sex matched mice were used between 8 to 11 weeks of age or at 20 weeks of age as stated in the text. All mice were housed and bred in specific pathogen-free conditions in the Animal Barrier Facility at the University of California San Francisco. Animal experiments were approved by the Institutional Animal Care and Use Committee of the University of California San Francisco.

### OVA model

The OVA model is outlined in Figure 1K. Mice were injected intraperitoneally with 50 µg OVA (Sigma-Aldrich) resuspended in 100 µl of sterile PBS and mixed with 100 µl of Imject alum (Thermo Fisher Scientific). After 7 days, the mice were challenged on three consecutive days by intranasal administration of 50 µg OVA resuspended in 20 µl of sterile PBS. Control mice received intranasal doses of 20 µl sterile PBS. 24 hours after the last intranasal challenge, the mice were anesthetized with isoflurane and 2 µg anti-CD45-BV510 antibody was injected by retroorbital route to label circulating blood cells. The mice were sacrificed 2-3 minutes later to collect BAL fluid and lung tissue. BAL was performed by 4 consecutive washes with 0.25 ml sterile PBS. BAL cells were incubated in ACK buffer (Lonza) for 2 min at RT to remove contaminating red blood cells, washed, resuspended in FACS buffer (2% FBS in PBS) and stored on ice until flow cytometry staining (described below). Airway inflammation was assessed by analysis of inflammatory cells in the BAL (eosinophils, neutrophils, alveolar macrophages, CD4^+^ T cells and CD8^+^ T cells), and frequency of Tregs was also measured in BAL. After BAL collection, lungs were PBS perfused by heart injection and left and right lung lobes were collected. Lung tissue was transferred to gentleMACS C-tubes (Miltenyi Biotec) containing 5 ml of freshly prepared lung digestion medium: 12.5 µg/ml DNase I (Sigma-Aldrich), 1.2 mg/ml Collagenase D (Roche) in RPMI 1640 (Fisher Scientific). The tissue was disrupted by running program m_lung_01 on the gentleMACS dissociator (Miltenyi Biotec) followed by 30 min incubation at 37°C with continuous shaking. Next, program m_lung_02 was used on the the gentleMACS dissociator and the tissue suspension was filtered through 70 µm Corning cell strainers (Fisher Scientific). Cells were incubated in ACK buffer for 2 min at RT, washed, resuspended in FACS buffer and stored on ice until flow cytometry staining of tissue-resident Tregs.

### In vivo proliferation by EdU

For EdU labeling of proliferating cells, 0.5 mg EdU in 200 µl sterile PBS was injected retroorbitally on three consecutive days before sacrifice, 24 hours after the last injection (Figure 2C). Thymus, spleen and inguinal lymph nodes were collected and cell suspensions were prepared by passing the tissues through 70 µm cell strainers. Spleen cells were incubated in ACK buffer for 2 min at RT. All cells were resuspended in FACS buffer and stored on ice until flow cytometry staining of EdU^+^ cells using the Click-iT Plus EdU Flow cytometry assay kit according to the manufacturer’s instructions (Thermo Fisher Scienticific).

### Treg transfer model

The colitis model by T cell transfer is outlined in Figure 3A. T cells were enriched from spleen and lymph nodes of CD45.1^+^ mice by negative selection using EasySep^TM^ Mouse CD4+ T cell isolation kit (STEMCELL Technologies), and FACS sorted as CD4^+^CD25^-^ CD44^lo^CD62L^hi^ cells. 400,000 naïve T cells were injected retroorbitally together with 150,000 Tregs, FACS sorted as CD4^+^FOXP3-GFP^+^ from spleen and lymph nodes. The Tregs were obtained from CD45.1^-^ miR-15/16f^l/fl^ *Cd4*-Cre mice or miR-15/16^fl/fl^ control mice. One group of mice received only 400,000 naïve T cells without Treg co-transfer. The weight of Tcrb KO recipient mice was recorded one time per week for 8 weeks. By the end of the model, colon, mesenteric lymph nodes and spleen were collected for flow cytometric analysis of Teff and Treg frequencies, and cytokine expression. Cell suspensions from spleen and lymph nodes were generated as described above. Colon was flushed with 10 ml of sterile PBS, cut in half longitudinally, then cut into small pieces and placed in a petri dish with 10 ml HEPES buffer: 15 mM HEPES (Fisher Scientific) in HBSS without calcium, magnesium, and phenol red (Fisher Scientific). The tissue pieces were gently agitated in the dish and washed in fresh HEPES buffer three times before being transferred to 10 ml 37°C digestion solution that consisted of RPMI-1640 supplemented with 2.5 mg/ml Collagenase D. Samples were incubated at 37°C with continuous shaking, and after 1h, the tissue suspension was filtered through 70 µm cell strainers. Remaining tissue pieces on the filter were placed in fresh digestion solution and incubated 1h at 37°C with continuous shaking. The procedure was repeated three times leaving no visible tissue pieces. Supernatants were centrifuged and cells were washed and resuspended in FACS buffer, pooling cells from all rounds of digestion.

### Flow cytometric analysis

Single cell suspensions from thymus, spleen, lymph nodes, colon, and lungs were transferred to 96-well conical bottom plates (Thermo Fisher Scientific) for antibody staining. Cells were washed one time in plain PBS followed by incubation for 15 min at 4°C in fixable viability dye at 1:2000 dilution in PBS. FACS buffer was used to wash, and the cells were blocked by adding anti-CD16/CD32 antibodies at 1:100 dilution in FACS buffer and incubated for 15 min at 4°C. Antibodies for surface stain was added at 1:100 dilution in blocking solution and incubated for 30 min at 4°C (dark). FACS buffer was used to wash the cells followed by fixation for 20 min at RT using Foxp3/Transcription Factor Staining Buffer set (Thermo Fisher Scientific). Surface-stained cells were washed, 5,000 AccuCount beads (Spherotech) were added, and samples were acquired on a BD LSRII flow cytometer (BD Biosciences). For samples that were stained intracellularly, fixation was followed by wash in permeabilization buffer (Foxp3/Transcription Factor Staining Buffer set). Intracellular antibodies were added at 1:100 dilution in permeabilization buffer and incubated for 1h at 4°C (dark). Samples were washed two times, 5,000 AccuCount beads were added, and samples were acquired on a BD LSRFortessa^TM^ cell analyzer (BD Biosciences). In experiments combining anti-FOXP3 antibody stain with endogenous FOXP3-YFP reporter, Foxp3/Transcription Factor Staining Buffer set was used as described above. Cells from lungs (OVA model) and mesenteric lymph nodes (colitis model) were re-stimulated ex vivo for cytokine staining and analysis by flow cytometry. In those experiments, cells were stimulated with 20 nM PMA and 1 µM ionomycin for 4h, where 5 µM brefeldin A was added during the final 2h. Following surface stain (described above), cells were fixed in 4% PFA for 8 min at RT. Ice cold plain PBS was added and cells were washed one time in cold plain PBS and one time in SAP permeabilization buffer (0.2% saponin in FACS buffer). Antibodies for intracellular cytokine stain was diluted at 1:20 concentration in SAP permeabilization buffer and added to the cells for 1h incubation at 4°C (dark). Cells were washed in FACS buffer and followed by acquisition on the BD LSRFortessa^TM^ cell analyzer. FlowJo V10 was used for the data analysis.

### In vitro Treg polarization

Naïve CD4^+^ T cells from spleen and lymph nodes were enriched by negative selection using EasySep^TM^ Mouse Naïve CD4+ T cell isolation kit (STEMCELL Technologies). 100,000 cells/well (50,000 CD45.1^-^ cells and 50,000 CD45.1^+^ cells) were added to 96-well plates (Costar 3370, Corning) coated with anti-CD3 (1 µg/ml) and anti-CD28 (0.5 µg/ml) antibodies. In IL-2 titration experiments, the cell culture medium was supplemented with 5 ng/ml recombinant human TGF-β (Peprotech), 10 µg/ml anti-IL-4 antibodies (clone 11B11), and 10 µg/ml anti-IFN-γ antibodies (clone XMG1.2) at seeding, and recombinant IL-2 (National Cancer Institute; concentrations indicated in figures) was added three days later when cells were moved to U-bottom 96-well plates with fresh culture medium. In TGF-β titration experiments, the cell culture medium was supplemented with TGF-β (as indicated in figures) at seeding along with anti-IL-4 and anti-IFN-γ antibodies at concentrations described above, and 20 units/ml IL-2 was added three days later. Cells were incubated at 37°C with 10% CO_2_ and iTregs were analyzed after 5 days in culture by flow cytometry.

### Ex vivo Treg cultures

CD4^+^ T cells were enriched from spleen and lymph nodes by magnetic separation and surface stained as described above. Cells were filtered and resuspended in sorting buffer (FACS buffer supplemented with 2 mM EDTA). Tregs were sorted as singlet live CD4^+^FOXP3-GFP^+^ cells on a BD FACSAria^TM^ II cell sorter. 100,000 Tregs/well were seeded in U-bottom 96-well plates, supplemented with recombinant IL-2 or IL-7 at concentrations indicated in figures, and incubated at 37°C with 10% CO_2_ ON. Cells were analyzed by flow cytometry the next day. For pSTAT5-staining, 500,000 CD4^+^ T cells enriched from spleen and lymph nodes were seeded in plain DMEM high glucose media and added to U-bottom 96-well plates. Cells were rested for 1h at 37°C with 10% CO_2_ before adding recombinant IL-2 or IL-7 in plain DMEM as indicated in figures. Cells were incubated 1h at 37°C with 10% CO_2_ and immediately fixed by adding 37°C 2% PFA followed by incubation for 10 min at RT. Cells were washed two times in FACS buffer then permeabilized with −20°C 100% methanol and stored ON in −20°C freezer. The next day, cells were washed two times in FACS buffer followed by rehydration for 20 min at RT in FACS buffer. Cells were washed and blocked as described above and incubated 30 min at RT (dark) with anti-STAT5 (pY694) antibodies (20 µl/test) and antibodies for surface stain (1:100 dilution). Cells were washed in FACS buffer and acquired on a BD LSRII flow cytometer where the endogenous FOXP3-YFP reporter was used to distinguish Tregs from other CD4^+^ T cells.

### In vitro suppression assay

CD45.1^-^ Tregs were FACS sorted as described above, and CD45.1^+^ Tcons were sorted as singlet live CD4^+^FOXP3-GFP^-^CD25^lo^ cells. Tregs and Tcons were CTV-labeled (5 µM in PBS) by incubation for 20 min at 37°C. Cells were washed and resuspended in complete cell medium (37). Splenocytes from Tcrb KO mice were resuspended in 50 µg/ml Mitomycin C (Roche) in PBS and incubated at 37°C for 1h (mixed repeatedly), followed by three repeated washes in plain PBS. Each well was supplemented with 5 µg/ml anti-CD3 (clone 145-2C11), 100,000 Mitomycin C-treated splenocytes and 50,000 CTV-labeled CD45.1^+^ Tcons. Tregs and Tcons were cultured at different seeding ratios, starting at 1:1 Treg:Tcon ratio, followed by 2-fold Treg dilution to have 2x, 4x and 8x more Tcons than Tregs. The cells were incubated for 72h at 37°C with 10% CO_2_ before flow cytometric analysis of proliferation by CTV dilution.

### Histology

Pancreas, the left lung lobe and the right liver lobe were placed in 10% formalin for 48h before transfer to PBS for storage. Skin samples were collected from the neck and placed in formalin for 5h then transferred to 30% sucrose for 48h followed by storage in PBS. Tissues were sectioned and H&E stained according to standard protocol.

### RNA sequencing and bioinformatic analysis

Tregs (bulk cKO, bulk WT and WT CD25^hi^ and CD25^lo^ bin) were sorted by FACS as described above using the endogenous FOXP3-YFP reporter. 340,000-1,200,000 cells were lysed in QIAzol (Qiagen) and total RNA was isolated using miRNeasy Mini Kit (Qiagen) with on-column DNase digestion. RNA quality and yield was determined on an Agilent 2100 Bioanalyzer (Agilent BioTek). cDNA was synthesized using the Nugen Universal Plus library kit (NuGen Technologies). The samples were sequenced using single-end 50 bp RNAseq on the Illumina HiSeq 4000 platform. Alignment was performed using STAR_2.7.2b against the Ensembl Human GRCm38.78 alignment genome. Differential expression was tested using DESeq2, and expression values were analyzed as log2 fold change of miR-15/16 deficient Tregs compared to miR-15/16 sufficient Tregs, or CD25^hi^ Tregs compared to CD25^lo^ Tregs.

Ingenuity Pathway Analysis (IPA; Qiagen) was used to identify upstream regulators of gene expression networks in Tregs dependent on CD25 or miR-15/16 expression. Venn diagrams were generated using Venny 2.1 (https://bioinfogp.cnb.csic.es/tools/venny/). Gene Set Enrichment Analysis (www.gsea-msigdb.org) was performed on 300 most upregulated or 300 most downregulated genes in Tregs with constitutive STAT5b expression compared to WT Tregs, and plotted against gene expression across miR-15/16 deficient or sufficient CD4^+^ T cells (22). Gene Set Enrichment Analysis was also performed on upregulated genes in miR-15/16 deficient Tregs compared to WT Tregs, and AHC >5 at miR-15/16 seed-matches in CD4^+^ T cells (22), plotted against the gene expression profile of eTreg or rTreg subsets (11). The cumulative distribution function plot was generated in R (www.r-project.org) to display the log2 fold change of DEGs (miR-15/16 deficient versus miR-15/16 sufficient Tregs) against the cumulative distribution of genes. The expression values from all genes lacking miR-15/16 3’UTR seed-match were plotted along with all genes containing a miR-15/16 3’UTR seed-match, a subset of all genes containing a miR-15/16 3’UTR seed-match with >5 AHC read mapping to the seed-match, and all genes predicted to be targeted by miR-15/16 by TargetScan (www.targetscan.org).

### Statistical analysis

Microsoft Excel and GraphPad Prism 9 were used for data analysis. In all figures, bar graphs are displayed with error bars showing standard deviation unless otherwise stated. Z-score was calculated from mean and standard deviation. *p < 0.05, **p < 0.01, ***p < 0.001, and ****p < 0.0001 for significance. ANOVA was used as statistical test of multiple groups, with appropriate post hoc testing as stated in the text. All data was assumed to be normally distributed.

### Data availability

Raw RNA-seq data and processed files reported in this paper will be available on GEO. miRNA expression data by RT-PCR in sorted human blood T cell populations was generated by Rossi *et al.* (NCBI GEO accession number GSE22880) (28). RNA-seq data of Tregs with constitutive STAT5b expression was generated by Chinen *et al*. (NCBI GEO accession number GSE84553) (6), and RNA-seq data of IL-7 stimulated mouse CD4^+^ T cells was generated by Villarino *et al.* (NCBI GEO accession number GSE77656) (52). Finally, RNA-seq data of miR-15/16 deficient and sufficient CD4^+^ T cells and AHC were generated by Gagnon *et al.* (NCBI GEO accession number GSE111568) (22), and RNA-seq data of TCF1^-^ LEF1^-^ Tregs was generated by Yang *et al.* (NCBI GEO accession number GSE117726) (11).

## Abbreviations

AHC: Ago2 high-throughput sequencing of RNAs isolated by crosslinking immunoprecipitation
BAL: Bronchoalveolar lavage
CTV: CellTrace^TM^ Violet
DEG: Differentially expressed genes
LN: Lymph node
MLN: Mesenteric lymph nodes
OVA: Ovalbumin
Tcon: Conventional T cell
Treg: Regulatory T cell
EdU: 5-ethynyl-2’-deoxyuridine

## Acknowledgements

We thank the UCSF SABRE Functional Genomics Core Facility for help with RNA sequencing; Mike Rosenblum and Jason Cyster for sharing mutant mice; and UCSF Parnassus Flow Cytometry CoLab (PFCC, (RRID:SCR_018206)) for use of their instruments.

This work was supported by the US NIH (HL107202, HL109102 and K24 HL137013), the Wenner-Gren Foundations, the Swedish Heart-Lung Foundation, the Sweden-America Foundation and the Foundation Blanceflor Boncompagni Ludovisi, née Bildt. The PFCC was supported in part by Grant NIH P30 DK063720 and by the NIH S10 Instrumentation Grant S10 1S10OD021822-01.

## Author contributions

K.J. and K.M.A. designed the research. K.J., J.D.G., S.Z., M.S.F., R.K., R.A.B., and H.P. performed the experiments. K.M.A., and P.G.W supervised the research. K.J., J.D.G., A.W.S., R.K., and P.G.W performed bioinformatic analysis under the advisement of K.M.A. K.J. and K.M.A. wrote the manuscript. All authors read and approved the final version of this manuscript.

