## Supplementary material for "An essential role for miR-15/16 in Treg suppression and restriction of proliferation": Supplementary information.pdf

Figure S1

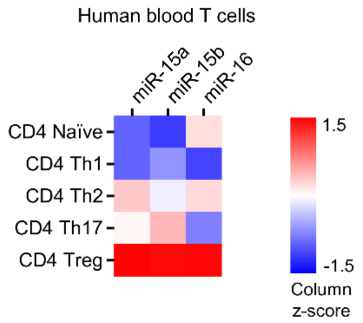


**Supplementary Figure S1. miR-15/16 expression in human T cell subsets.**
qPCR of miRNAs purified from $CD4^+$ T cell subsets in peripheral blood of healthy donors (Rossi *et al.* (1); GSE22880). Flow cytometry was used for isolation of T cell subsets; CD4 Naïve $CD4^+CCR7^+CD45RA^+CD45RO^-$; CD4 Th1 $CD4^+CXCR3^+CCR6^-CD161^-$; CD4 Th2 $CD4^+CRTH2^+CXCR3^-$; CD4 Th17 $CD4^+CCR6^+CD161^+CXCR3^-$; CD4 Treg $CD4^+CD127^{lo}CD25^+$. N=3-6/group.

Figure S2

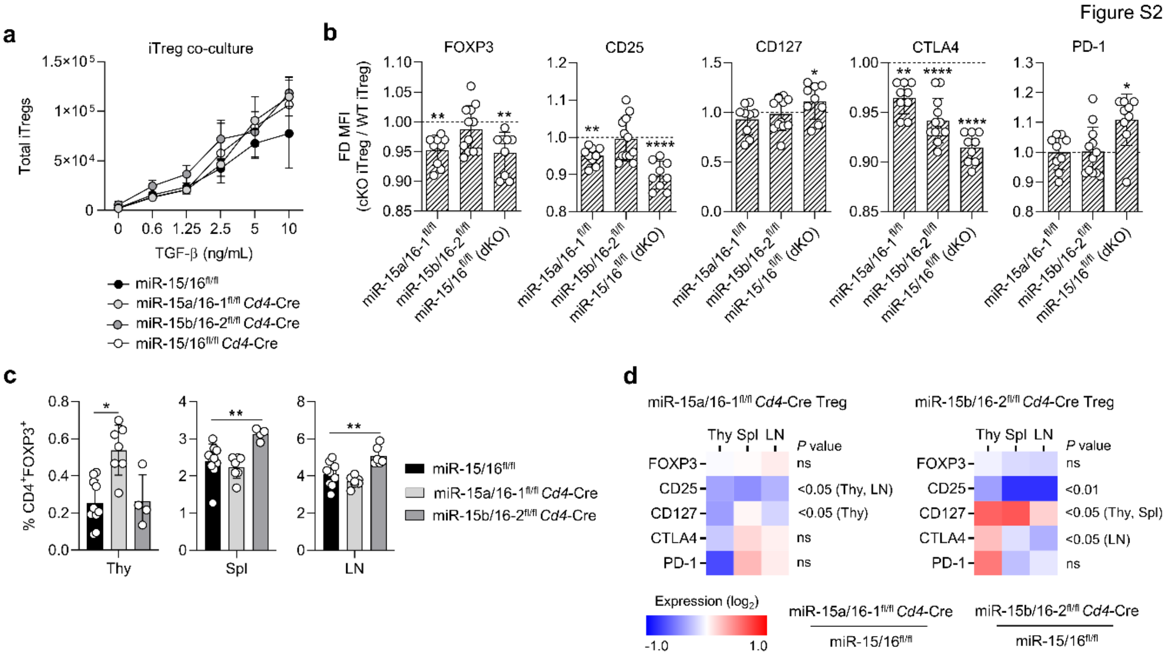


**Supplementary Figure S2. miR-15/16 single cluster expression modulate Treg phenotype.**

**A:** Total number of induced Tregs (iTregs) after 5 days in co-culture under Treg polarizing conditions assessed by FOXP3 expression by flow cytometry. **B:** Protein expression by median fluorescent intensity (MFI) compared to co-cultured WT control iTregs. **C:** Frequency of Tregs among all T cells of miR-15a/16-1$^{fl/fl}$ *Cd4*-Cre mice, miR-15b/16-2$^{fl/fl}$ *Cd4*-Cre mice and miR-15a/16-1$^{fl/fl}$ control mice. **D:** Fold change of MFI by flow cytometry of indicated proteins in Tregs of miR-15a/16-1$^{fl/fl}$ *Cd4*-Cre mice (left) and miR-15b/16-2$^{fl/fl}$ *Cd4*-Cre mice (right) from three tissues (change in knockout from miR-15/16$^{fl/fl}$ WT control).

Figure S3

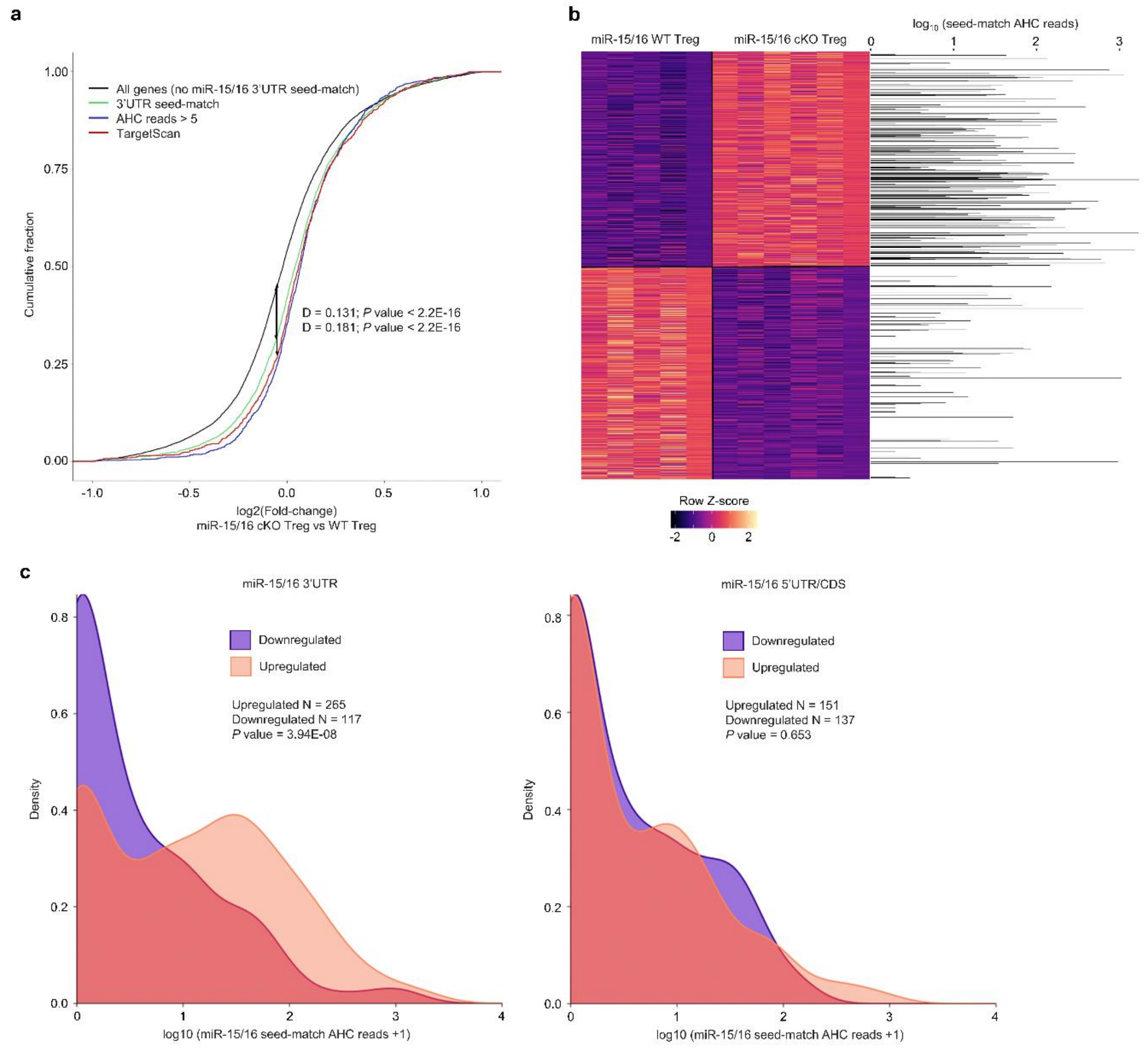


**Supplementary Figure S3. miR-15/16 bind and regulate direct target RNAs in Tregs.** **A:** Cumulative density plot depicting global expression by RNA sequencing as a ratio of the fold change between miR-15/16$^{fl/fl}$ *Foxp3*$^{Cre}$ ('miR-15/16 cKO Treg'; n = 6 biological replicates) and miR-15/16$^{wt/wt}$ *Foxp3*$^{Cre}$ ('miR-15/16 WT Treg'; n = 5 biological replicates) Tregs for all genes without a 7-mer or 8-mer miR-15/16 3'UTR seed match (black), genes with a 7-mer or 8- mer miR-15/16 3'UTR seed match (green), genes with a 7-mer or 8-mer miR-15/16 3'UTR seed match and AHC read depth >5 (blue), and genes classified as targets of miR-15/16 by TargetScan 7.0 (red) (AHC reads represent the combined depth of n = 10 independent immunoprecipitations). **B:** Heatmap of genes with a P-value <0.05 plotted alongside a bar graph of AHC read depth at miR-15/16 seed matches for each gene at which they occur. **C:** Comparison of AHC reads between genes that are downregulated and upregulated (P<0.05) in miR-15/16$^{fl/fl}$ *Foxp3*$^{Cre}$ Tregs among genes with seed matches in the 3'UTR (left) or 5'UTR/CDS (right) (Mann-Whitney U test). AHC data was generated by Gagnon *et al.*(2); GSE111568.
